# Engineering substrate selectivity in the human sodium/iodide symporter (NIS)

**DOI:** 10.1101/2025.10.07.681042

**Authors:** Alejandro Llorente-Esteban, Haswitha Sabbineni, Kendra Hoffsmith, Rían W. Manville, David López-González, Andrea Reyna-Neyra, J. Alfonso Leyva, Geoffrey W. Abbott, Mario A. Bianchet, Nancy Carrasco

**Affiliations:** Department of Molecular Physiology and Biophysics, Vanderbilt University, Nashville, TN 37232, USA; Department of Physiology and Biophysics, School of Medicine, University of California, Irvine, CA 92697, USA; Pontificia Universidad Javeriana, Bogotá, Colombia; Department of Biophysics and Biophysical Chemistry, Johns Hopkins University School of Medicine, Baltimore, MD 21205, USA

## Abstract

Iodide (I^-^) uptake mediated by the Na⁺/I^-^ symporter (NIS) is the first step in the biosynthesis of the thyroid hormones, of which I^-^ is an essential constituent. NIS couples the inward transport of I^-^ against its electrochemical gradient to the inward translocation of Na^+^ down its electrochemical gradient. NIS also transports oxyanions (XO_4_^-^s), such as perrhenate (ReO_4_^-^) and the environmental pollutant perchlorate (ClO_4_^-^). Furthermore, NIS is the basis for radioiodide (¹³¹I^-^) therapy for thyroid cancer (administered after thyroidectomy), the most effective targeted internal radiation cancer therapy available. ^131^I^-^ selectively targets remnant malignant cells and metastases expressing NIS, causing only minor side effects. There is great interest in expressing NIS exogenously, by gene transfer, in extrathyroidal cancers to render them susceptible to destruction by ^131^I^-^. This approach, however, would also harm patients’ thyroids. Therefore, a strategy is needed for killing non-thyroidal cancer cells exogenously expressing NIS while protecting the thyroid. Addressing this need, we present here an engineered double mutant, L253P/V254F (PF)-NIS, which selectively transports XO_4_^-^s but not I^-^. We used cryo-EM to determine the structure of PF-NIS with ReO_4_^-^ and Na^+^ ions bound to it at a 2.58 Å resolution, and showed that high concentrations of non-radioactive I^-^ protect WT-NIS-expressing cells from radioactive ^186^ReO_4_^-^, whereas PF-NIS-expressing cells are killed. Thus, PF-NIS could potentially be used, together with ^186/188^ReO_4_^-^ and non-radioactive I^-^, to treat non-thyroidal cancers while safeguarding the thyroid. This study establishes a framework for developing therapies using NIS molecules engineered to have selective substrate specificities to extend the clinical use of NIS beyond thyroid cancer.

## Introduction

The sodium (**N**a^+^)/**I**^-^ **s**ymporter (**NIS**)^1,2^ is the key plasma membrane protein that actively transports I^-^ into the thyroid^2–4^. NIS couples the inward transport of I^-^ against its electrochemical gradient to the inward translocation of Na^+^ down its electrochemical gradient, with an electrogenic 2 Na^+^ : 1 I^-^ stoichiometry^2^. NIS-mediated I^-^ transport into the thyroid is the first step in the biosynthesis of the thyroid hormones (THs) T_3_ and T_4_, of which I^-^ is an essential constituent. The THs are crucial for the maturation of the central nervous system, skeletal muscle, and respiratory system in intrauterine and early life, and are master regulators of intermediary metabolism in virtually all tissues^2^. NIS transports several other substrates besides I^-^, including oxyanions (XO_4_^-^s), such as pertechnetate (^99m^TcO_4_^-^), perrhenate (ReO_4_^-^), and the environmental pollutant perchlorate (ClO_4_^-^). XO_4_^-^ transport by NIS is electroneutral, with a 1 Na^+^ : 1 XO_4_^-^ stoichiometry^5^; thus, NIS transports different substrates with different stoichiometries. NIS is the molecule that makes it possible to treat differentiated thyroid cancer with radioiodide (^131^I^-^) (administered after thyroidectomy), the most effective targeted internal radiation cancer therapy available. ^131^I^-^ selectively targets remnant malignant cells and metastases expressing NIS, resulting in only minor side effects. This treatment has been used since 1946 ^6^, 50 years before our group first isolated the cDNA encoding NIS, which enabled us to identify the transporter and characterize it at the molecular level^3^.

Since then, there has been great interest in expressing NIS exogenously by gene transfer in extrathyroidal cancers, thus rendering them susceptible to destruction by ^131^I^-^ ^7–10^. However, if NIS is exogenously expressed in extrathyroidal cancer cells and ^131^I^-^ is administered to destroy them, the radioisotope will also damage the thyroid. Therefore, an effective strategy is needed for protecting the thyroid. To this end, we sought to engineer a NIS molecule specifically designed to transport XO_4_^-^s but not I^-^. With such a molecule expressed exogenously in non-thyroidal cancer cells, patients could be treated with radioactive perrhenate (^186/188^ReO_4_^-^) instead of ^131^I^-^, and the thyroid could be shielded with excess non-radioactive I^-^.

Two key developments paved the way for the present study. First, over the years, several mutations in NIS have been identified in patients with I^-^ deficiency disorders (IDDs), the study of which has yielded valuable insights into the protein’s folding, plasma membrane targeting, and activity^11–28^. Second, we have determined three structures of NIS by cryo-EM: apo-NIS, NIS with 2 Na^+^ and 1 I^-^ bound to it, and NIS with 1 Na^+^ and 1 ReO_4_^-^ bound to it ^29^. The structures, along with extensive mutagenesis and detailed functional studies, have provided a wealth of mechanistic information on NIS. In addition, a recent report of a NIS mutation in transmembrane segment (TMS) 7 (G250V)^22^ in patients with congenital hypothyroidism led us to investigate that particular region of the protein.

Here we show that, by replacing only two amino acids in TMS 7, we engineered the double mutant L253P/V254F NIS (PF-NIS), which has the desired substrate selectivity: it transports XO_4_^-^s but virtually no I^-^. Moreover, the mechanism by which PF-NIS transports XO_4_^-^s is electrogenic, unlike that of WT-NIS, which transports XO_4_^-^s electroneutrally. We have used cryo-EM to determine the structure of PF-NIS with ReO_4_^-^ and Na^+^ ions bound to it, at a resolution of 2.58 Å. Remarkably, we showed in clonogenic survival assays that PF-NIS-expressing cells are killed by ^186^ReO_4_^-^ even in the presence of large amounts of non-radioactive I^-^, at concentrations high enough to protect WT-NIS-expressing cells. Our findings demonstrate that NIS molecules with selective specificity can be engineered to extend the clinical use of NIS beyond thyroid cancer.

## Results

### Glycine 250 is key for NIS activity

The G250V substitution in NIS (Fig. 1a) has recently been reported in patients with congenital hypothyroidism^22^. However, it was not stated how the mutation affected NIS activity. To address this question, we generated an MDCK cell line that stably expresses G250V-NIS. The typical electrophoretic pattern of WT-NIS in Western blots consists of a prominent broad band at ∼100 kDa (fully glycosylated NIS) and a minor band at ∼65 kDa (partially glycosylated NIS) (Fig. 1b, lanes 1 and 3). In contrast, the pattern for G250V-NIS featured a predominant ∼65 kDa band and a barely detected ∼100 kDa band, suggesting that most G250V-NIS molecules were not fully glycosylated (Fig. 1b, lanes 5 and 7). Indeed, endoglycosidase H (Endo H) treatment caused the ∼65 kDa polypeptide to migrate faster (at ∼50 kDa), but had no effect on the ∼100 kDa polypeptide (Fig. 1b, lanes 4 and 8). For its part, peptide-N-glycosidase F (PNGase F) digested both polypeptides (∼65 and 100 kDa) (Fig. 1b, lanes 2 and 6), indicating that a small fraction of the G250V-NIS molecules proceed beyond the medial Golgi to acquire complex N-glycans. Consistent with these findings, our flow cytometry experiments showed that only 35% of the G250V-NIS molecules were at the plasma membrane, compared to 86% of WT-NIS molecules (Extended Data Fig. 1). G250V-NIS was not only targeted less efficiently to the plasma membrane, but was also inactive (Fig. 1c, inset). Out of nine NIS mutants with amino acid substitutions at position 250, only G250P-NIS transported I^-^, and only 16% as much as WT-NIS (Fig. 1c). In contrast, G250P-NIS transported 50% as much ^99m^TcO_4_^-^ as WT-NIS (Fig. 1d). Interestingly, though, this may be because proline, though rigid, imparts to this region of the protein the “right kind” of stiffness, thereby mimicking the structural role of glycine just enough for the protein to retain partial activity. This region is where the α-helix transitions into a π-helix [residues 247 to 251, as determined using the Define Secondary Structure of Proteins (DSSP) program] (Fig. 1a). π-helices are flexible structural elements often found near functional sites.

**Fig. 1.**
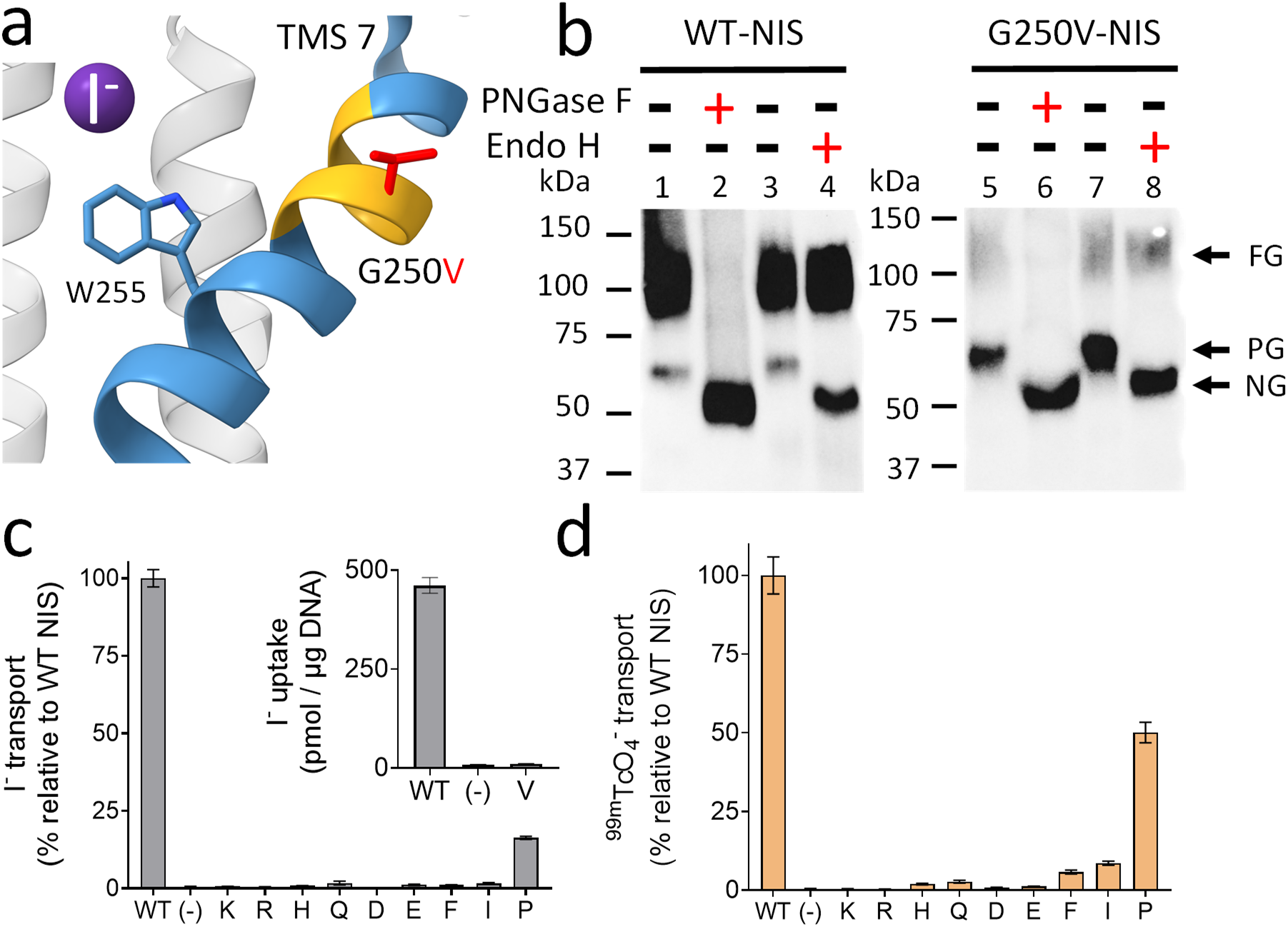
| NIS function requires an amino acid at position 250 that renders the π-helix of TMS 7 flexible. **a,** View of TMS 7 in the WT-NIS model (PDB: 7UV0), as determined by the cryo-EM density map with I^-^ (purple sphere) bound to it. TMS 7 and residue W255 are highlighted in blue. The most likely rotamer of V250 is shown in red, in between the π-helix residues (yellow). **b**, Western blot of MDCK cells expressing WT- or G250V-NIS. We loaded 5 μg of membrane preparation for WT-NIS and 7.5 μg for G250V-NIS. Proteins loaded in lanes 2 and 6 were digested with PNGase F, and those loaded in lanes 4 and 8 were digested with Endo H. Proteins loaded in lanes 1, 3, 5, and 7 were not digested (FG: fully glycosylated NIS; PG: partially glycosylated NIS; NG: non-glycosylated NIS). **c,** Steady-state I^-^ transport by nine mutant NIS proteins with substitutions at position 250. Inset: transport by WT- or G250V-NIS; non-NIS-expressing MDCK cells were used as a negative control (-). **d,** Steady-state ^99m^TcO_4_^-^ transport by the mutant NIS proteins as in (c). Transport data are averages of three biological replicates (n = 3); each experiment was performed in triplicate, and results are presented as mean ± s.e.m.

### Altering the substrate selectivity of NIS

Because G250V-NIS partially discriminates between I^-^ and XO_4_^-^s, and because position 250 is close to W255 (Fig. 1a), which is crucial for NIS activity^16^, we investigated the significance of L253 and V254 for NIS function. Whereas the side chain of L253 points away from the protein’s core (Fig. 2a), the side chain of V254 faces the substrate binding cavity, although without interacting directly with the substrates (Fig. 2a)^29^. We introduced amino acid substitutions at positions 253 and 254 and showed that the expression levels of the resulting mutants varied depending on the substitution (Extended Data Fig. 2a, b). Only L253R-NIS was not fully glycosylated. Most 253 mutants translocated approximately as much I^-^ as WT-NIS, whereas charged residues at this position markedly impaired I^-^ transport. Interestingly, the L253P and L253H substitutions reduced I^-^ transport only moderately (40% and 60% as much I^-^ as WT-NIS, respectively) (Fig. 2b). Most 253 mutants transported ^99m^TcO_4_^-^ better than I^-^ (Fig. 2c). Remarkably, L253P- and L253H-NIS, which transported less I^-^ than WT-NIS, transported approximately as much ^99m^TcO_4_^-^ as WT-NIS (Fig. 2b, c).

**Fig. 2.**
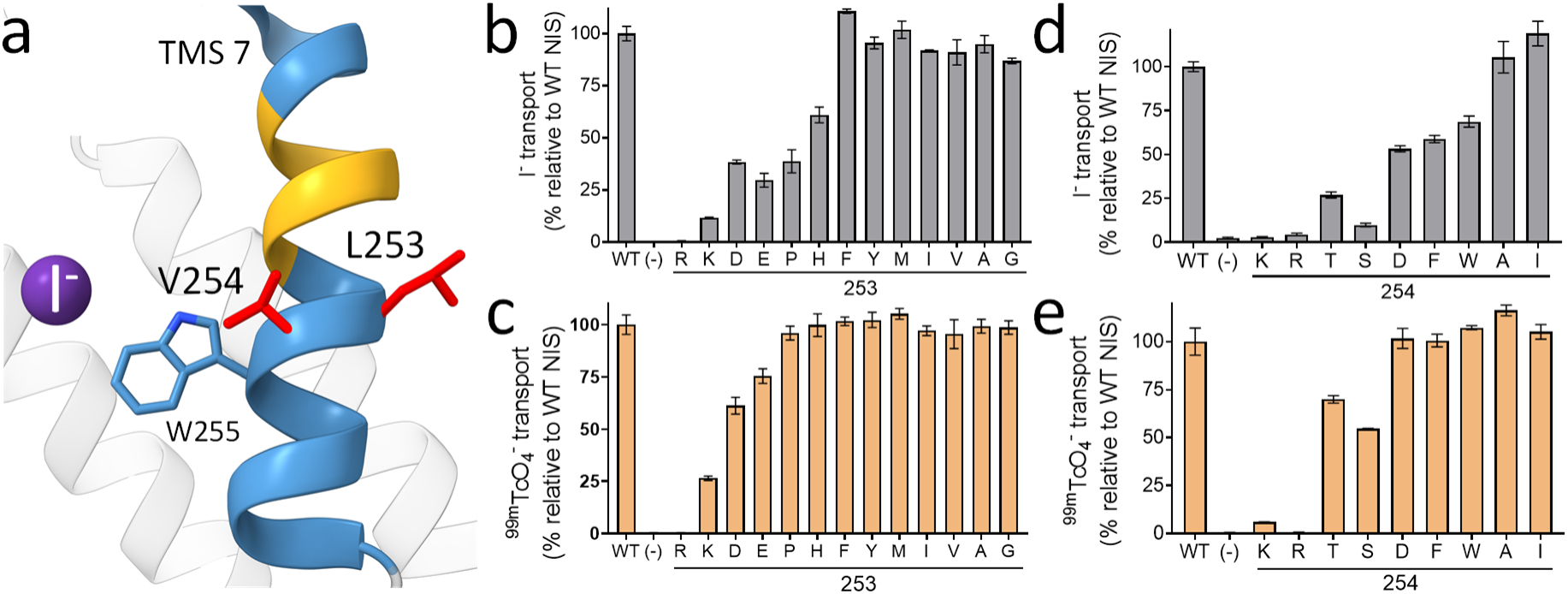
| Amino acid substitutions at positions 253 and 254 cause NIS to partially discriminate between its substrates. **a,** View of TMS 7 from a different angle in the WT-NIS model (PDB: 7UV0). TMS 7 and the residue W255 are highlighted in blue, and L253 and V254 in red; the latter are adjacent to the π-helix (yellow). **b, c, d,** and **e**, Steady-state ^125^I^-^ (**b** and **d**) and ^99m^TcO_4_^-^ (**c** and **e**) transport by NIS mutant proteins with amino acid substitutions at position 253 (**b** and **c**) or 254 (**d** and **e**). For panels (b), (c), (d), and (e), data were background-subtracted and normalized with respect to transport by WT-NIS. Data in (b) and (c) are representative of two biological replicates performed in triplicate (n = 6), and data in (d) and (e) are representative of three biological replicates performed in triplicate (n = 9). Data are presented as mean ± SD.

Overall, substitutions at position 254 had a greater impact on NIS function than those at position 253. Most of the 254 mutants also transported ^99m^TcO_4_^-^ better than I^-^ (Fig. 2d, e). We found that—like L253P-and L253H-NIS—V254D-, V254W-, and V254F-NIS transported less I^-^ than WT-NIS, but approximately the same amount of ^99m^TcO_4_^-^ as WT-NIS (Fig. 2d, e). That certain substitutions adjacent to the π-helix of TMS 7 caused NIS to partially discriminate between I^-^ and XO_4_^-^s was quite interesting, especially because some of those mutants transported XO_4_^-^s at near-WT levels, but transported less I^-^ than WT-NIS. Notably, L253H- and L253P-NIS transported 1.6 and 2.5 times as much ^99m^TcO_4_^-^ as I^-^, respectively (Extended Data Fig. 2c), and V254F-, V254W-NIS translocated between 1.5 to 1.9 times as much ^99m^TcO_4_^-^ as I^-^ (Extended Data Fig. 2d). In summary, certain substitutions at positions 253 and 254 clearly alter the substrate specificity of NIS.

### Engineering an oxyanion-selective NIS mutant for cancer therapy

To extend NIS-mediated targeted internal radiation therapy to non-thyroidal cancers while protecting the thyroid, a NIS molecule must be engineered that selectively transports XO_4_^-^s but no I^-^, as it would allow the use of radioactive ^186^ReO_4_^-^ to kill the cancer cells and non-radioactive I^-^ to shield the thyroid (Extended Data Fig. 3). Having shown that some single substitutions yield mutants with substrate specificities different from those of WT-NIS, we generated the double mutants L253P-V254F (PF-NIS) and L253H-V254F (HF-NIS) to investigate whether the effects of the double substitutions would be synergistic. The levels of expression of PF- and HF-NIS in MDCK cells were similar to those of WT-NIS (Fig. 3a); however, PF- and HF-NIS barely transported I^-^ (less than 7% and 2% of WT-NIS levels, respectively) (Fig. 3b). On the other hand, HF-NIS transported 33% as much ReO_4_^-^ as WT-NIS and, remarkably, PF-NIS transported as much ReO_4_^-^ as WT-NIS (Fig. 3c). Therefore, PF-NIS is a molecule with the substrate specificity we were seeking.

**Fig. 3.**
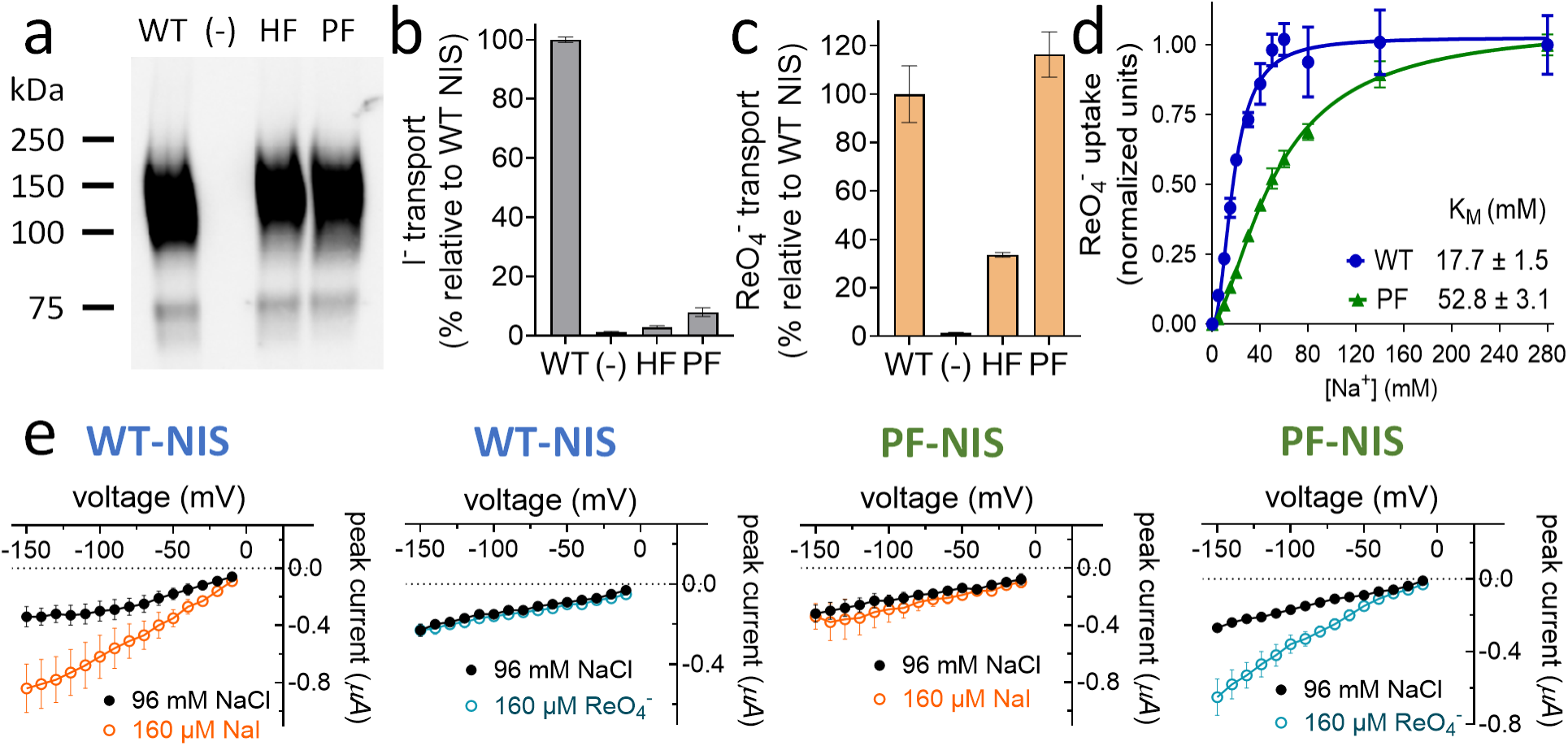
| The double mutant PF-NIS fully discriminates between I^-^ and ReO_4_^-^. **a,** Western blot of MDCK cells expressing the NIS double mutants L253P & V254F (PF) and L253H & V254F (HF). WT-NIS was used as a positive control, and non-NIS-expressing MDCK cells as a negative control (-). 7 µg of membrane preparations were loaded for all samples. **b** and **c**, Steady-state I^-^ (b) and ^186^ReO_4_^-^ (c) transport by the NIS double mutants. Results shown in (b) and (c) are representative of three biological replicates performed in triplicate (n = 9); data were normalized with respect to transport by WT-NIS; error bars represent SD. **d,** Initial rates of 10 μM ReO_4_^-^ transport as a function of increasing concentrations of Na^+^ by WT- and PF-NIS. Values represent averages of the results from three different experiments, each carried out in duplicate (n ≥ 6). Error bars represent s.e.m. **e,** Current–voltage relationships recorded by TEVC in oocytes upon addition of 160 μM I^-^ (orange; n = 5 for WT-NIS, n = 7 for PF-NIS) or 160 μM ReO_4_^-^ (blue; n = 5 for WT-NIS, n = 6 for PF-NIS). Background currents were recorded in the absence of substrates at 96 mM NaCl (black). For (d) and (e), error bars represent s.e.m.

The K_M_ of PF-NIS for ReO_4_^-^ (K_M_ = 6.5 ± 0.7 μM) was comparable to that of WT-NIS (K_M_ = 1.4 ± 0.2 μM) (Extended Data Fig. 4), and the K_M_ of PF-NIS for Na^+^ (K_M_ = 52.8 mM) was higher than that of WT-NIS (K_M_ = 17.7 mM) (Fig. 3d). Unexpectedly, PF-NIS exhibited cooperativity for Na^+^ when transporting ReO_4_^-^, suggesting that PF-NIS-mediated XO_4_^-^ transport is electrogenic (Fig. 3d). This switch in stoichiometry was confirmed by electrophysiological experiments in *Xenopus laevis* oocytes, which showed that positive inward currents were elicited by ReO_4_^-^ only in PF-NIS-expressing but not in WT-NIS-expressing oocytes (Fig. 3e, blue lines). As expected, currents were elicited by I^-^ only in WT-NIS-expressing^16,30^, but not in PF-NIS-expressing oocytes (Fig. 3e, orange lines). In sum, the PF-NIS molecule we engineered not only transports exclusively XO_4_^-^s, but transports them electrogenically.

### Structural insights into the transport mechanism of PF-NIS

To understand at the atomic level how PF-NIS discriminates between I^-^ and XO_4_^-^s, we solubilized and purified PF-NIS with ReO_4_^-^ bound to it (Extended Data Fig.5), and collected and processed cryo-EM data (Extended Data Fig. 6). The density map of the PF-NIS monomer reaches a local resolution of ∼2.4 Å for most of the protein, except for TMS 1 (local resolution ∼ 3.5 Å), which is highly mobile and located at the periphery of NIS (Fig. 4a and Extended Data Fig. 7), resulting in a global resolution of 2.58 Å (Extended Data Fig. 6g and Extended Data Table 1).

**Fig. 4.**
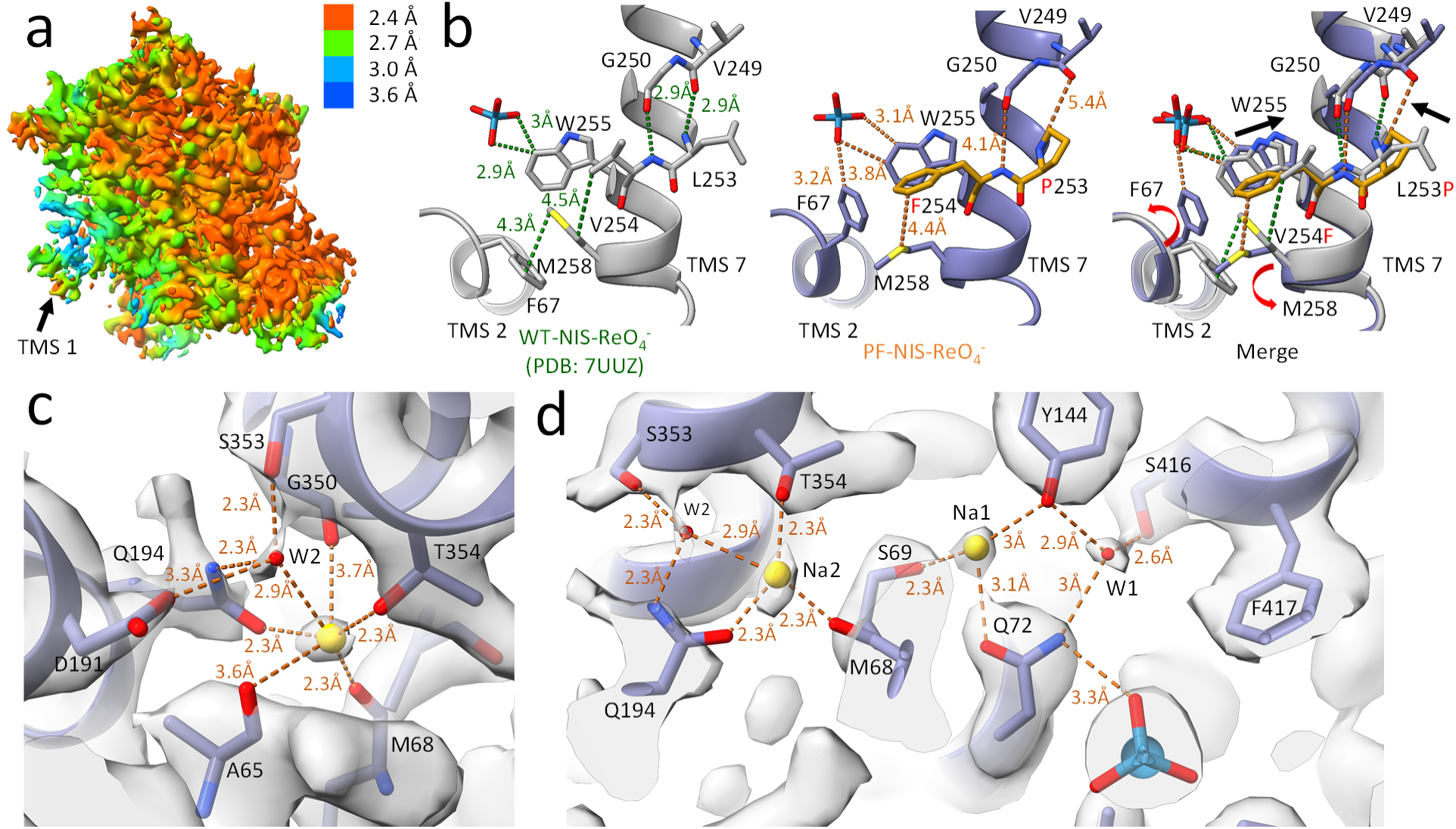
| Structure of PF-NIS. **a,** Density map colored according to local resolution. **b,** Conformational changes induced by L253P and V254F in PF-NIS (middle panel) and WT-NIS (left panel). Black arrows in the merged figure (right panel) indicate the main movements. **c,** Newly identified density at the Na2 site. **d,** Na1 and W1 sites showing communication between them via Q72 in TMS 2, and between Na1 and Na2 via S69 and M68 in the unwound region of TMS 2.

When we compared the atomic model of PF-NIS bound to ReO_4_^-^ with our previously reported WT-NIS-ReO_4_^-^ structure^29^ (RMSD = 0.59 Å for 492 Cα atoms), we found that ReO_4_^-^ binds to both PF-NIS and WT-NIS at the same site (Extended Data Fig. 8a). However, the L253P and V254F substitutions introduce key conformational changes at the ReO_4_^-^ binding site. First, the V254F mutation places the aromatic ring of F254 perpendicular to W255 (Fig. 4b, middle panel), thereby restricting the side-chain mobility of W255. This effect is clearly illustrated by the side-chain dihedral angles (χ₁, χ₂) derived from 6.8 µs of aggregated molecular dynamics (MD) simulations for WT-NIS–ReO_4_^-^ and 7.3 µs for PF-NIS–ReO_4_^-^: in WT-NIS, W255 explores a broader range of conformational states (Extended Data Fig. 9a); in PF-NIS, by contrast, the rotations of the W255 side chain are constrained to a considerable degree (Extended Data Fig. 9b). Second, in the PF-NIS structure, the Pro introduced at position 253 prevents an H-bond from forming with the main-chain carbonyl of V249, pushing the helical turns in which they are located apart and introducing a kink into the helix (Fig. 4b, middle and right panels). This kink, in turn, breaks the H-bond between the carbonyl of G250 and the amide of F254; the distance is 2.9 Å in WT-NIS but, importantly, 4.1 Å in PF-NIS (Fig. 4b); interestingly, G250 is also the residue that was mutated in the patient. The loss of these bonds causes a bend around residues 249–251, slightly retracting W255 from the substrate-binding cavity and further reducing the mobility of W255, and increasing its distance from ReO_4_^-^ from 2.9 Å to 3.8 Å (Fig. 4b, right panel). Despite the retraction of W255, the side chain of F254 narrows the cavity from ∼83.5 Å³ in WT-NIS- ReO_4_^-^ (PDB: 7UUZ) to ∼72.7 Å³ in PF-NIS. This contraction likely explains why PF-NIS has a reduced apparent affinity for ReO_4_^-^. The overall result is a tighter packing around the XO_4_^-^. The steric hindrance from F254 and helix destabilization by P253 render the contracted cavity more rigid, consistent with MD simulations showing that the side chain of W255 is less mobile in PF-NIS than in WT-NIS (Extended Data Fig. 9b). Moreover, the bulkier side chain of F254 causes M258 to shift toward the translocation pathway (Fig. 4b), creating steric hindrance that limits the “aromatic swing” of F67. In the WT-NIS structures, F67 acts as a gating (*gauche⁻*) residue at the base of the substrate cavity, facilitating the release of the substrates into the cytosol (Fig. 4b, left panel)^29^. In PF-NIS, F67 adopts a new conformation, with its side-chain in the *trans* rotamer (χ_1_ ≈ 180; χ₂ ≈ 80°) (Extended Data Fig. 9c, d), pointing towards the anion binding site and coordinating ReO_4_^-^ through hydrophobic interactions. This results in a more occluded NIS conformation (Fig. 4b, middle panel). However, in PF-NIS, F67 also displays an unstable rotamer, consistent with its weaker electron density (Extended Data Fig. 7). Comparative MD simulations of WT-NIS–ReO_4_^-^ and PF-NIS–ReO_4_^-^ show that the mutations reduce the probability of sampling the gating *gauche⁻* rotamer, χ ≈ 300°, χ₂ ≈ 100° (Extended Data Fig. 9c, d). W255 and F67 function as gates that enable NIS substrates to be transported from the ion-binding cavity into the cytosol. In WT-NIS, the conformational flexibility of W255 and F67 allows the protein to transition efficiently from the inwardly occluded to the inwardly open state; in PF-NIS, on the other hand, the mobility of those residues is restricted. Consistent with that, PF-NIS requires a greater driving force to complete its transport cycle: its Na^+^/ReO_4_^-^ stoichiometry is not 1:1, as in WT-NIS, but 2:1, as shown in our transport and electrophysiology experiments (Fig. 3 d, e).

Aside from TMS 1 at the periphery of the protein, and the side chains of F67 and M68—which lie in the highly mobile unwound region of TMS 2, which is indirectly impacted by the V254F substitution (Fig. 4b)— the high-quality cryo-EM density in PF-NIS enabled us to structurally trace the protein unambiguously (Extended Data Fig. 7). We can now discern with greater clarity an isolated tetrahedral density within a cavity formed by residues Y118, Y137, A140, S257, V261, N262, S424, and L427 (Extended Data Fig. 8d). This density could be due to a physiological oxyanion (e.g., sulfate, phosphate, bicarbonate, or nitrate). It is unlikely to be due to the ReO_4_⁻ added to the sample, as a similar density is present in the structure of WT-NIS with I⁻ bound to it and likely in the apo-NIS structure (ReO_4_⁻ was not added to these structures and is not a physiological anion). That said, the issue merits further investigation, given the considerable resolution differences between the PF-NIS and WT-NIS structures.

In our new, higher-resolution PF-NIS structure, we clearly observe a density at a site equivalent to the canonical Na2 site, first identified in the bacterial leucine transporter (LeuT) of *Aquifex aeolicus*^31^. In WT- NIS, a Na⁺ interaction at this site has been proposed to drive I^-^ transport^11,32,33^. In PF-NIS, Na⁺ is coordinated at this Na2 site by the carbonyls of M68 and in close proximity to A65 in TMS 2, the carbonyl of G350 in TMS 9, and the hydroxyl group of T354 (equivalent to S355 in TMS 8 of LeuT) (Fig. 4c). Additionally, a water molecule (W2) stabilized by S353 (equivalent to T354 in TMS 8 of LeuT) also coordinates Na⁺ (Fig. 4c). In PF-NIS, contrary to the situation observed with LeuT, the flexible side chain of Q194 in TMS 6 also participates in coordinating Na^+^ at the Na2 site through an ionic bond (Fig. 4c). The side-chain oxygen of Q194 sits 2.3 Å from the Na⁺, and the water molecule W2 coordinating the Na⁺ ion is stabilized by interactions with D191, Q194, and S353, forming a tetrahedral arrangement around W2 (Fig. 4c). Of note, we previously proposed—on the basis of the results of MD simulations—that, in WT-NIS, D191 and Q194 interact with Na⁺, and we demonstrated that both residues are important for NIS function^33^.

We suggest that the movement of F67, triggered by the V254F substitution, changes the positions of key backbone residues in the unwound region of TMS 2—including M68—to favor Na⁺ binding at this site, yielding a near-complete octahedral coordination characteristic of metal-binding sites.

In the PF-NIS structure, we observed two additional densities at positions corresponding to those previously dubbed the “Na1” and “Na2” sites in WT-NIS (Extended Data Fig. 8e). At the time, we attributed these densities to Na⁺ ions, a hypothesis supported by the results of a coulombic potential analysis^29^. Importantly, the site previously labeled “Na2” in WT-NIS does not correspond to the canonical Na2 site first found in LeuT^31^ that we also see now in PF-NIS. To avoid confusion and keep to the standard terminology in the field, we now refer to what was previously called the Na2 site in WT-NIS as Na1, to what was previously called the Na1 site in WT-NIS as W1 (see below), and to the site analogous to LeuT’s Na2 site as Na2 (Fig. 4d and Extended Data Fig. 8e).

The density at what we are now calling the Na1 site remains well-defined in PF-NIS, coordinated by the side chains of Q72 (3.1 Å), S69 (1.9 Å), and Y144 (3 Å) (Fig. 4d). In PF-NIS, Na1 is further stabilized by the main-chain carbonyl of F67 at 2.5 Å (Extended Data Fig. 8b). The side chain of this residue could plausibly also contribute a cation–π interaction with the bound ion, though the distances observed are on the longer side for that type of coordination (Extended Data Fig. 8b). These interactions between this Na⁺ and F67 are not observed in WT-NIS—but in PF-NIS, as previously mentioned, the side chain of F67 adopts a high-energy conformation in the mutant protein owing to the steric hindrance induced by the V254F substitution (Fig. 4b), which also positions F67 closer to the Na1 site.

The density at what we are now calling the W1 site is no weaker than those at the Na1 or Na2 sites. The density at the W1 site has a tetrahedral coordination provided by the OH groups of Y144 (2.9 Å) and S416 (2.6 Å), and the amide group of Q72 (3 Å) (Fig. 4d), consistent with an assignment of this density to a water molecule. Furthermore, this water molecule could be additionally coordinated by the Na^+^ ion assigned with confidence to the Na1 site, when it transitions to occupy an alternative position at a Na1’ site, which is closer to the ReO_4_^-^ binding site (Extended Data Fig. 8c). Na^+^ at Na1’ would be coordinated by cation–π interactions with the aromatic ring of F417 and the water molecule at W1. In addition, Na⁺ would interact with ReO₄⁻, stabilizing the oxyanion and neutralizing its negative electrostatic potential (discussion and Extended Data Fig. 8c).

To further investigate the densities observed at the Na1, Na1’, and Na2 sites in PF-NIS, we performed MD simulations using a system with PF-NIS embedded in a mammalian phospholipid membrane at near-physiological conditions, including ion gradients and a membrane potential of -70 mV (see Materials and Methods) and periodic boundary conditions. Extended Data Figure 9e shows the aggregated 7.3 µs time series of the ions’ *z* coordinates in PF-NIS simulations loaded with ReO_4_^-^ and three cations placed at the cryo-EM–observed sites. Across 16 replicas, the system rapidly converges to a two-cation state because the Na^+^ originally placed at the Na1’ site escapes into the extracellular milieu. The other 2 Na^+^ ions are transported into the cytosol; Na2 stays bound for longer, consistent with its better coordination (Fig. 4c). ReO_4_^-^ remains bound in most trajectories, but comes close to being transported in 9 out of 16 replicas, in which the Na^+^ ions actually are translocated into the cytosol (Extended Data Fig. 9e).

### ^186^ReO_4_^-^ kills PF-NIS-expressing cells even in the presence of I^-^

Given the remarkable transport properties of PF-NIS (Fig. 3b-e) and our insights into its structure [which are consistent with its electrogenic transport of ReO_4_^-^ (Fig. 4c, d)], we investigated the feasibility of combining PF-NIS and radioactive ^186^ReO_4_^-^ to target and destroy cells—while protecting cells expressing WT-NIS by saturating the endogenous transporter with non-radioactive I^-^ (Extended Data Fig. 3). First, we performed experiments to infer the apparent affinity of PF-NIS for I^-^, using ^99m^TcO_4_^-^ as a tracer and increasing concentrations of I^-^. The K_I_ of PF-NIS for I^-^ (K_I_ = 215.1 μM) was 10 times higher than that of WT-NIS (K_I_ = 21.9 μM) (Fig. 5a). Notably, ^99m^TcO_4_^-^ transport by PF-NIS was not completely inhibited by I^-^, even at concentrations as high as 800 μM, which is hundreds of times the submicromolar physiological concentration of I^-^ ^34^. These results show that, at a concentration of 100 μM I^-^, 86% of the ^99m^TcO_4_^-^ transport mediated by WT-NIS is inhibited, whereas, remarkably, PF-NIS continues transporting as much ^99m^TcO_4_^-^ as it does in the absence of 100 μM non-radioactive I^-^ (Fig. 5a, red dotted line).

**Fig. 5.**
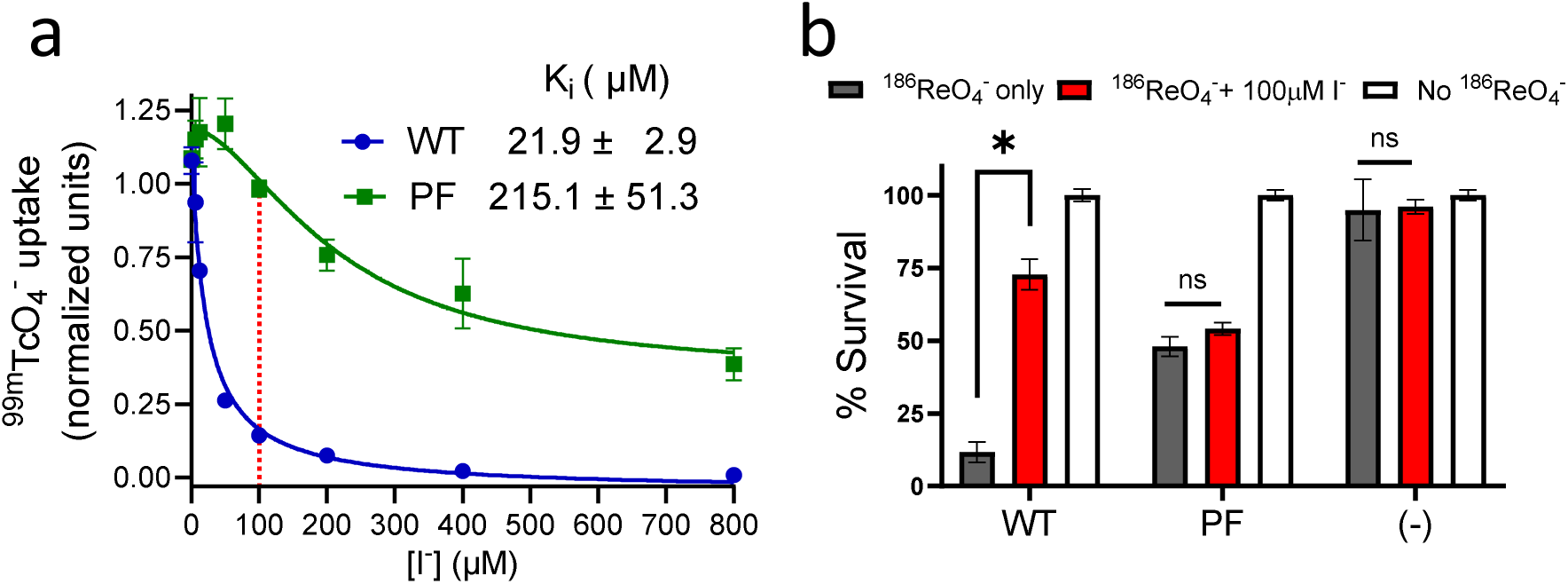
| Saturating concentrations of non-radioactive I^-^ protect cells expressing WT-NIS from ^186^ReO_4_^-^, but not those expressing PF-NIS. **a,** Initial rates of ^99m^TcO_4_^-^ transport as a function of increasing concentrations of non-radioactive I^-^ by WT- and PF-NIS. Representative experiment of two biological replicates carried out in triplicate (n = 6). Error bars represent SD. **b,** Clonogenic survival experiment in MDCK cells expressing WT- or PF-NIS, as well as control non-NIS-expressing cells (-). Cells were exposed to three conditions: no radioactivity (empty bars; n = 12) or 0.033 mCi/ml of radioactive ^186^ReO_4_^-^ (grey bars; n = 18), or ^186^ReO_4_^-^ with 100 μM I^-^ (red bars; n = 18). Data represent the mean ± s.e.m. of three biological replicates. Differences between groups were analyzed using unpaired Welch’s t-tests.

As a proof of concept then, we performed a clonogenic survival assay by exposing MDCK cells expressing WT-NIS or PF-NIS to three different conditions: HBSS without radioactive ReO_4_^-^, as a control; HBSS/^186^ReO_4_^-^ (33 µCi/ml); and HBSS/^186^ReO_4_^-^ (33 µCi/ml) with 100 μM I^-^, to protect cells expressing WT-NIS but not those expressing PF-NIS (based on the results in Fig. 5a). Of the cells expressing WT-NIS, >80% died (∼20% survived) when incubated with ^186^ReO_4_^-^ alone; by contrast, with ^186^ReO_4_^-^ plus 100 μM non-radioactive I^-^, only ∼25% died, because I^-^ saturated WT-NIS, thereby protecting the remaining ∼75% surviving cells from ^186^ReO_4_^-^ (Fig. 5b). Crucially, the percentage of cells expressing PF-NIS that died when incubated with ^186^ReO_4_^-^ and 100 μM I^-^ (∼50%) was *<u>no different</u>* from the percentage of cells that died when they were *<u>only</u>* incubated with ^186^ReO_4_^-^ (Fig. 5b). These data show that, *in vitro*, 100 μM I^-^ protects cells expressing WT-NIS from the effects of radioactive ^186^ReO_4_^-^, but has no effect on cells expressing the engineered protein PF-NIS (Fig. 5b). All control non-NIS-expressing cells survived, indicating that cells were only killed by ^186^ReO_4_^-^ when it was actively transported into the cytoplasm by NIS (Fig. 5b).

## Discussion

The high specificity and effectiveness of NIS-mediated targeted internal radiation therapy for thyroid cancer, along with its low incidence of even relatively minor side effects, are major reasons why there is great interest in developing strategies for using this treatment modality against other cancers. The principle at the heart of this therapy is remarkably elegant: the radionuclide targets NIS-expressing malignant cells, while largely sparing all other cells. Moreover, a key factor that plays a role in the efficacy of this treatment is the “cross-fire effect,” i.e., the fact that even non-NIS-expressing tumoral cells that are within a short distance of NIS-expressing cells are also killed because of the range of the emitted β radiation^7,35,36^. The prospect of extending the benefits of targeted internal radiation therapy for thyroid cancer to non-thyroidal cancers began to come into view over two decades ago, when our group first isolated the cDNA encoding NIS^3^. At that point, it became possible to carry out pre-clinical studies ectopically expressing NIS by gene transfer in cancer cells that do not express it endogenously, thus rendering them susceptible to NIS-mediated targeted internal radiation treatment, with either ^131^I^-^ ^37,38^ or ^186/188^ReO_4_^-^ ^39,40^. Furthermore, ^186^ReO_4_^-^ and ^188^ReO_4_^-^ are more energetic and have shorter half-lives than ^131^I^-^(^186^ReO_4_^-^: *E* = 0.35 MeV, t = 3.8 days; ^188^ReO_4_^-^: *E* = 0.764 MeV, t = 17 hrs; ^131^I^-^: *E* = 0.134 MeV, t = 8 days)^36^, and, although ReO_4_^-^ is actively transported by NIS into the thyroid, it is advantageously not stored in the gland, as it is not covalently incorporated into tyrosyl residues in the thyroid hormone precursor thyroglobulin—unlike iodine (oxidized I⁻), which is.

Although these advantages may increase the efficacy of therapy with ^186/188^ReO_4_^-^, using these radionuclides instead of ^131^I^-^ still would not protect the thyroid, as WT-NIS would also transport ^186/188^ReO_4_^-^.

Several pre-clinical studies investigating the effects of exogenous NIS-mediated targeted internal radiation therapy in rodent models of aggressive cancers—including those of the prostate, pancreas, and liver, as well as glioblastoma—have shown effective tumor shrinkage or growth delay^8–10,39,40^. In these studies, NIS was delivered to the tumors using viral vectors^8,39,40^, nanoparticles^9,10^, or mesenchymal stem cells^41,42^. Still, to effectively administer this therapy to patients, the optimal delivery vector and the appropriate tumor-specific promoter or specific receptor ligand would need to be established for each type of cancer. In addition, a major hurdle remains: finding a way to protect the patients’ thyroids from the radionuclide. One way to tackle this problem is by administering THs, which will downregulate NIS expression in the thyroid, thus markedly decreasing (but not completely preventing) radionuclide uptake by the gland, as we previously demonstrated in humans^43^. But in addition to not completely preventing radionuclide uptake by the thyroid, this strategy has a second major drawback: it causes hyperthyroidism, which makes it unfeasible.

That being so, we generated a NIS molecule, PF-NIS, which selectively transports XO_4_^-^s but translocates virtually no I^-^ (Fig. 3b and c), and it mediates XO_4_^-^ transport electrogenically—and using not only the Na^+^ electrochemical gradient but also the membrane potential as a driving force (Fig. 3d and e). Expressing PF-NIS in non-thyroidal cancer cells would make it possible to treat patients by targeting their malignant cells with ^186/188^ReO_4_^-^, while the concomitantly administered non-radioactive I^-^ would saturate thyroidal endogenous WT-NIS, thereby selectively preventing accumulation of the radionuclide in the gland (Extended Data Fig. 3). Because of the aforementioned “cross-fire effect,” the treatment with ^186/188^ReO_4_^-^ would kill not only PF-NIS-expressing but also neighboring non-PF-NIS-expressing malignant cells.

Because PF-NIS has such interesting properties, we determined the structure of this mutant with ReO_4_^-^ bound to it by cryo-EM, at a resolution of 2.58 Å (Extended Data Fig. 6g). We propose that PF-NIS does not transport I^-^ because, even if I^-^ is one of the largest monatomic anions (with an ionic radius of 2.2 Å), it is still smaller than ReO_4_^-^ (ionic radius = 2.60 Å), and thus has fewer polar contacts than the oxyanion within the more constricted and rigid substrate cavity observed in the PF-NIS structure (Fig. 4b and Extended Data Fig. 6g). Therefore, the oxygen atoms in the ReO_4_^-^ molecule are likely to form stronger polar interactions with the protein, whereas the weaker electrostatic interactions that I⁻ can establish may be insufficient to facilitate effective translocation through the pathway altered by the double amino acid substitution. Thus, the restricted mobility of W255 and F67 and decreased cavity size appear to be sufficient to prevent PF-NIS from transporting I^-^ while still allowing it to translocate ReO_4_^-^. Furthermore, the hydrogen bond between residues 254 and 250 explains why the patient’s G250V mutation impairs I^-^ transport: the increased rigidity in the π-helix likely reduces the flexibility required for the conformational changes that NIS must undergo in order to be able to transport its substrates.

The PF-NIS structure shows two densities that we assigned to Na⁺ ions at the Na1 and Na2 sites, and one density that we assigned to a water molecule, at the W1 site (Fig. 4d). The near-octahedral coordination of the Na⁺ ion at the Na2 site (Fig. 4c) and the longer dwelling times observed in MD simulations for the Na⁺ ion at this site (Extended Data Fig. 9e) suggest that the Na2 site corresponds to the high-affinity Na⁺ binding site in NIS, while the Na1 site binds Na⁺ with lower affinity, as suggested by the reduced coordination number. Furthermore, the structural relationship between the Na2 and Na1 sites explains the cooperativity observed between the two Na⁺ ions transported by NIS^44,45^. The Na⁺ at the Na2 site is coordinated by the main-chain carbonyl of M68, whereas the Na⁺ at the Na1 site is coordinated by the hydroxyl group of S69. Notably, M68 and S69 are consecutive residues located in the highly dynamic unwound region of TMS 2, which enables the Na1 and Na2 sites to communicate directly (Fig. 4d).

Examining more closely the region between W1 and F417 allowed us to propose an alternative interpretation of the density map according to which the lower-affinity Na⁺ ion toggles between the Na1 site—which stabilizes the unwound region of TMS 2—and, perhaps, a Na1’ site, which is closer to the negatively electrostatic environment induced by ReO_4_^-^, and hence would help ReO_4_^-^ and Na^+^ stabilize each other (Extended Data Fig. 8c). The density corresponding to the F417 side chain is crescent-shaped, and near it—at low thresholds—there is a weak density between the W1 site and F417, which we dubbed Na1’ (Extended Data Fig. 8c).

When Na⁺ is located at the Na1’ site, it may be stabilized at this position by a different rotamer of F417 that makes possible a cation–π interaction, and by the water molecule at W1 (Extended Data Fig. 8c). Thus, this alternative explanation (prompted by what the electron density map revealed about how NIS interacts with its substrates) suggested that the Na1 and Na1’ sites are partially occupied by a single Na⁺ ion. In other words, this weak density could represent a site that (along with the Na1 site) is partially occupied by Na^+^, and stabilized by the anion, the water molecule at W1, and a possible cation-π interaction with an alternative rotamer of F417 (Extended Data Fig. 8c). In this scenario, a single Na⁺ bound at the Na1 site— and toggling between the Na1 and Na1’ sites—would be transported together with the Na⁺ ion at the Na2 site.

Armed with these new insights into how PF-NIS distinguishes between I^-^ and ReO_4_^-^, we now turn to its potential therapeutic applications. Consistent with widely accepted guidelines for protecting the thyroid with non-radioactive I^-^ in the event of a nuclear accident ^46,47^—and as hypothesized—we demonstrated in a clonogenic survival assay that non-radioactive I^-^ effectively protects WT-NIS-expressing cells from the deleterious effects of radioactive ^186^ReO_4_^-^ (Fig. 5b). In sharp contrast, we showed—as a proof of principle— that PF-NIS-expressing cells are killed by the radionuclide even in the presence of saturating concentrations (100 µM) of non-radioactive I^-^ (Fig. 5b). Crucially, the amount of radioactive ¹⁸⁶ReO_4_^-^ used in our study was considerably lower (20-30 times less) than the dose employed in previous *in vitro* survival experiments using WT-NIS^39,40^. Furthermore, we used ^186^Re instead of the more energetic ^188^Re which would have been more cytotoxic. Clearly, the treatment conditions will need to be further optimized in future *in vivo* studies.

In conclusion, although the engineered substrate specificity of PF-NIS is very promising—as is the fact that using it in conjunction with radioactive ^186/188^ReO_4_^-^ and non-radioactive I^-^ is likely to cause few side effects—animal studies are needed to determine the efficacy of this therapy. Still, this study shows unequivocally that long-term basic structural, mechanistic, and functional studies of membrane transport proteins, such as NIS, can enable scientists to design new molecules for optimizing therapeutic interventions with the potential to benefit a large number of patients.

## Materials and methods

### Cell Culture

We employed Madin-Darby canine kidney (MDCK-II) cells that stably express either the human WT-NIS or its variants with specific amino acid substitutions at positions 250, 253, and 254. The MDCK cells were cultured in high glucose Dulbecco’s Modified Eagle Medium (DMEM). This medium was supplemented with 10% fetal bovine serum, 100 units/ml penicillin-streptomycin and 2 mM L-glutamine, all materials obtained from Life Technologies, Inc. All cell cultures were incubated at 37°C in a humidified atmosphere containing 5% CO_2_.

We purified PF-NIS using 293F cells that stably express the mutant protein following lentiviral transduction. The PF amino acid substitution was introduced into a rat NIS cDNA background engineered to be unglycosylated, by mutating the relevant asparagine residues to glutamine. Our previous work established that this unglycosylated construct functions identically to the glycosylated version, showing no defects in transport, expression or trafficking to the plasma membrane^16,17,19,29,48^. Moreover, the construct carries an HA tag at its extracellular N-terminus and a streptavidin-binding protein tag at its intracellular C-terminus. For cell culture, 293F cells were maintained in 293 SFM II medium (Gibco™ Cat. No. 11686029, ThermoFisher SCIENTIFIC) supplemented with 4 mM L-glutamine (Life Technologies, Inc.). Cultures were grown in 1 L flasks to a density of 300,000–3,000,000 cells/mL, incubated at 37°C in a humidified atmosphere containing 8% CO₂, and agitated at 120 RPM.

### Transport experiments

Transport experiments were conducted using the method previously documented^19,49^. Briefly, MDCK cells were plated on 24-well plates one day prior to experimentation. At the end of the experiment, the accumulated isotope was extracted using ice-cold ethanol and quantified using a Cobra-II Auto-gamma Counter from Packard Bioscience. For all functional experiments, the amount of uptake was standardized to the amount of DNA per well. ^125^I^-^ transport experiments were performed at steady state at a specific activity of 1.5 µCi ^125^I^-^/ml, incubating the cells for 1 hour at 37°C. ^99m^TcO_4_^-^ transport experiments were conducted identically. ^125^I^-^ was obtained from Perkin-Elmer, Boston, MA, USA, and ^99m^TcO_4_^-^ from the Vanderbilt University Institute of Imaging Science, Nashville, TN, USA.

The transport kinetics for ReO_4_^-^ were conducted with a varied range of ReO_4_^-^ concentrations from 0.675 µM to 40 µM, maintaining the Na^+^ concentration constant at 140 mM. The specific activity of ^186^ReO_4_^-^ was 100 µCi/µmol, obtained from the University of Missouri MURR, Columbia, MO, USA. The background was acquired and subtracted using non-expressing NIS MDCK cells. The inhibition transport kinetics of ^99m^TcO_4_^-^ were performed with 1.5 µCi/ml of ^99m^TcO_4_^-^ as a tracer, along with increasing concentrations of non-radioactive I^-^ ranging from 6.25 to 800 µM with a constant Na^+^ concentration of 140 mM.

Na^+^-dependent kinetics of ReO_4_^-^ transport were executed with ^186^ReO_4_^-^ and increasing Na^+^ concentrations from 0 to 280 mM. Here, osmolarity was kept uniform (280 mM) with choline chloride. Non-radioactive ReO_4_^-^ was used at the indicated concentrations in the corresponding figure legend. The background was acquired and subtracted from the values obtained of cells expressing NIS when these were incubated with cocktails containing 0 mM Na^+^ (so NIS cannot mediate transport), as well as from negative controls using cells that do not express NIS. All kinetics were conducted at initial transport rates (4 minutes). Transport data were plotted and analyzed using Prism 10 software (GraphPad Software Inc.). Initial rate data for ReO_4_^-^ and ^99m^TcO_4_^-^ transport were fitted using nonlinear regression analysis with equations selected based on the experimental setup: ReO_4_^-^ transport as a function of increasing Na⁺ concentrations was analyzed using an allosteric sigmoidal equation; ReO_4_^-^ transport as a function of increasing ReO_4_^-^ concentrations was analyzed using the Michaelis–Menten equation; and ^186^ReO_4_^-^ transport as a function of increasing I^-^ concentrations was analyzed using the [inhibitor] vs. response – variable slope equation.

### Preparation of cRNA and Injection into *Xenopus laevis* oocytes

cDNAs of WT and mutant NIS were first linearized and then transcribed *in vitro* using the T7 polymerase mMessage kit (Thermo Fisher Scientific). The concentration of the resulting cRNA was determined by spectrophotometric analysis. Stage V and VI *Xenopus laevis* oocytes (Xenoocyte, Dexter, MI) were then defolliculated and injected with 10–20 ng of cRNA. The injected oocytes were kept at 16 °C in Barth’s saline solution (Xenoocyte) containing penicillin and streptomycin, with the medium refreshed daily, for a period of five days before conducting two-electrode voltage-clamp (TEVC) recordings.

### Two-Electrode Voltage Clamp (TEVC)

Five days after cRNA injection, TEVC recordings were conducted at room temperature (20–22 °C) using an OC-725C amplifier (Warner Instruments) and pClamp10 software (Molecular Devices). Oocytes were positioned in a small-volume bath (Warner) and observed under a dissection microscope. All reagents were sourced from Sigma. The bath solution composition was (in mM): 96 NaCl, 4 KCl, 1 MgCl₂, 1 CaCl₂, and 10 HEPES (pH 7.6). Stock solutions of ReO_4_^-^ and I^-^ were prepared at 1 M and diluted to working concentrations on the day of each experiment. Anions were introduced into the oocyte bath via gravity perfusion at a steady rate of 1 ml/min for three minutes before data collection. Recording pipettes had a resistance of 1–2 MΩ when filled with 3 M KCl. Currents were measured in response to step pulses from - 150 mV to -10 mV in 10-mV increments, with a holding potential of -50 mV, to generate current-voltage curves. Data were analyzed using GraphPad Prism 10.

### Flow Cytometry and Cell Sorting

Immunofluorescent procedures were employed for cell staining as outlined in previously established methodologies^16,19^. Initially, cells fixed with paraformaldehyde were exposed to a solution containing 0.2% BSA in PBS, supplemented with a monoclonal anti-HA IgG from rats (clone 3F10, 3 nM), produced by Roche Applied Science. Note that the HA tag engineered at the extracellularly-facing N-terminus, does not interfere with NIS function or trafficking to the plasma membrane^16,17,19,48^. For studying total NIS expression under permeabilized conditions, the cells were also incubated with saponin (an additional 0.2%), and the anti-HA IgG from rats (clone 3F10, 3 nM) was added to the solution. Following rinsing, the cells were subsequently incubated with a goat antibody conjugated to phycoerythrin that recognizes rat antigens (anti-rat antibody acquired from Life Technologies). Cell fluorescence was assessed utilizing a mass cytometry technology instrument: the MACSQuant flow cytometer by Miltenyi Biotec. The 293 cells expressing PF-NIS were sorted, to obtain a population in which over 99% of the cells expressed the engineered mutant, using a high-speed cell sorter (5-laser FACSAria III). This sorting was conducted under sterile conditions, without permeabilization and using the same IgGs than for analytical flow cytometry experiments. Analysis of the obtained data was completed using FlowJo software developed by Tree Star.

### Immunoblot

Homemade SDS/7.5% acrylamide PAGE TGX stain-free gels (Bio-Rad) were used for our electrophoretic separation. We extracted membrane preparations from stable cell lines expressing either WT-NIS or its mutant variants. All samples were accurately diluted in a ratio of 1:2 with the loading buffer, followed by heat treatment at 37°C for 30 minutes to ensure sample readiness for electrophoresis. The separated proteins were subsequently transferred to a PVDF membrane using a standard electroblotting procedure as delineated in prior studies^50^. All immunoblot analysis procedures were performed by incubating with an anti-C-terminal human or rat NIS antibody at 1:3000 dilution and a horseradish peroxidase-conjugated sheep anti-rabbit IgG (Chemicon International) at 1:5000 dilution. These incubations were executed either at room temperature for one hour or overnight at 4°C to enable antibody-antigen binding. A chemiluminescent image of the bound antibodies was obtained using a ChemiDoc MP Imaging System (Bio-Rad) to visualize the polypeptides. Equal amounts of protein loaded and transferred to the PVDF membrane were assured by taking advantage of the Bio-Rad stain-free technology.

### Deglycosylation experiments

Membrane preparations derived from cells expressing either WT- human NIS or the NIS variant G250V were subjected to deglycosylation processes with either PNGase F or Endo H (New England Biolabs, Beverly, MA). In the initial phase of the experiment, 40 µg of the prepared membranes were denatured for 30 minutes at 37°C. For the PNGase F deglycosylation, 10 µg of the prepared membranes and 1% (vol/vol) Nonidet P-40 were combined with 1000 U PNGase F, and the reaction was maintained for 2 hours at 37°C. Similarly, for the Endo H deglycosylation, 10 µg of the membrane preparations were mixed with the NEB buffer for Endo H and 500 U of Endo H enzyme, with the reaction conditions being maintained at 37°C over a period of 2 hours. Corresponding control samples were incubated in identical buffers, albeit without enzymes. All samples were then analyzed using immunoblot, performed as outlined above.

### Protein purification

The purification began with 293F cells with stable expression of PF-NIS. These cells were collected and suspended in a buffered solution containing 75 mM Tris-HCl (pH 8.0), 350 mM NaCl, and protease inhibitors (Roche). Cell disruption was achieved mechanically using an Emulsiflex C3 homogenizer (Avestin), and the debris were removed by spinning the mixture at 10,000g for 10 minutes. The membrane-rich supernatant was then subjected to ultracentrifugation at 250,000g for two hours to isolate the membrane fraction. To solubilize these membranes, they were incubated at 4°C for two hours in a buffer supplemented with 0.5% lauryl maltose neopentyl glycol (LMNG) and 0.5% glyco-diosgenin (GDN). PF-NIS was subsequently purified through Strep-Tactin affinity chromatography, followed by size exclusion chromatography using a Superdex 200 Increase column equilibrated with 350 mM NaCl, 75 mM Tris-HCl (pH 8.0), 0.005% LMNG, and 0.005% GDN. The fractions containing the purified protein were collected and prepared for electron microscopy imaging.

### Cryo-EM Sample Preparation and Data Processing

Freshly glow-discharged Quantifoil 1.2/1.3 grids (Cu 300 mesh, Electron Microscopy Sciences) were utilized for cryo-EM sample preparation. Following purification, the isolated NIS molecules were incubated with 1 mM ReO_4_^-^. Subsequently, 2.5 μl of protein at a concentration of 0.5 mg/ml was applied to the grids. The grids were blotted for 6 seconds at 100% humidity and 7°C before being vitrified in liquid ethane using a Vitrobot Mark IV (Thermo Fisher Scientific).

The cryo-EM dataset for PF-NIS was acquired at the Center for Structural Biology Cryo-Electron Microscopy Facility at Vanderbilt University using a ThermoFisher Titan Krios G4 electron microscope operating at 300 kV and equipped with a Gatan K3 BioQuantum direct detector. Images were captured at a pixel size of 0.647 Å with a total accumulated dose of 55 e^-^/Å^2^. Drift correction of the video stacks was performed using Patch Motion Correction, followed by Patch CTF Estimation in CryoSPARC v4.5.3 ^51^.

Out of 33,939 total images, 29,245 exposure-curated images were accepted. Initial particle picking with CryoSPARC’s Blob Picker selected 11,890 particles on a cropped dataset, which underwent two rounds of 2D classification. The best classes were used to perform a template picker job across the full dataset, resulting in 9,754,870 picked particles, which were subsequently extracted using a 360 px box size. Following three rounds of 2D classification, 816,391 particles were selected to create an Ab-initio models.

Subsequently, one round of heterogeneous refinement, additional 2D classification, and another round of heterogeneous refinement reduced the dataset to 462,777 selected particles with a resolution of 2.96 Å. Particles were split by exposure group, with their beam shift corrected using Global CTF refinement, accounting for beam tilt and trefoil aberrations. A final reference-based motion correction yielded a 2.80 Å average resolution for the NIS dimer using non-uniform refinement.

Given that both monomers in the non-physiological dimer are identical, symmetry expansion was applied. The density of one of the monomers was extracted, and local refinement was performed using a mask covering the unsubtracted monomer. The final PF-NIS atomic model was generated using Coot^52^, starting with the WT-NIS structure in the presence of ReO₄⁻ (PDB: 7UUZ), introducing the corresponding mutations, and refining with the Phenix Real-Space Refine tool^53^ (Extended Data Table 1).

### Structural Analysis

Structural representation of NIS in the presence of its substrates were retrieved from our previous published structures (PDB: 7UV0; 7UUZ)^29^, and our newly PF-NIS structural data, and subsequently examined using the UCSF ChimeraX software^54^. This enabled the visualization of amino acid positions 250, 253, and 254. The “rotamers” tool (Dunbrack library) available within the UCSF ChimeraX software was deployed to generate the most plausible rotamer for G250V over the atomic model of WT-NIS in the presence of I^-^ (PDB: 7UV0). All representative pictures were also created using the same software.

### Clonogenic Survival Assay

MDCK cells, stably transfected with either human WT- or PF-NIS, were sorted to ensure that over 90% of the cells were expressing the protein. The cells were subsequently cultivated in addition within a 25-cm^2^ flask. Prior to incubation, cells were rinsed twice with HBSS, and then exposed to: 3 ml of HBSS containing 33.3 μCi/ml of radioactive ^186^ReO_4_^-^; or 33.3 μCi/ml of radioactive ^186^ReO_4_^-^ and 100 μM I^-^; or HBSS that lacked the radioisotope ^186^ReO_4_^-^, for a period of 2 hours. The termination of the reaction involved the removal of the medium that contained the radioisotope and washing the cells twice using HBSS. The cells were then trypsinized and counted, following which they were divided and seeded in six replicates at a density of 1000 cells per well.

For culturing, the cells were placed in DMEM high glucose medium, dispersed across 6-well plates. Culturing continued for seven days post-seeding, after which the cells were fixed using 4% paraformaldehyde and stained with a solution combining 0.5% crystal violet and 20% methanol. A GelCount™ colony counter (Oxford Optronix) was used to identify and quantify macroscopic colonies, with the criteria being over 50 cells per colony. The survival rate percentage was evaluated as the number of colonies (per well) that survived to the exposure to ^186^ReO_4_^-^ treatments, versus the number of colonies resulting when the cells were treated without the radioisotope. The clonogenic survival assay was plotted and analyzed using the software Prism 10 developed by GraphPad.

### Statistics analysis

For the clonogenic survival assay, data were analyzed using GraphPad Prism 10. Multiple unpaired t-tests were performed using the Welch t-test, which does not assume equal variances between groups. This method was chosen to account for potential unequal variances across groups. A p-value < 0.05 was considered statistically significant for each comparison. Results are presented as mean ± standard error of the mean (s.e.m).

### MD Simulations of PF-NIS

A near-physiological condition simulation system consisting of one NIS molecule embedded in the central membrane with the asymmetric mammalian lipid composition, subject to the physiological ions gradient between two assembled non-connected solvent chambers separated by a simpler auxiliary membrane and to the difference of potential across the membrane. The phosphatidylcholine (POPC) and cholesterol lipids form an auxiliary membrane, forming two separated chambers on both sides of the main membrane by the action of the system periodicity in the membrane normal (z). The two separated chambers were filled with water and ions (Na^+^, K^+^, and Cl^-^) at concentrations representatives of the extra- and intracellular media at either side of the main membrane, respectively, to generate the physiological ions gradient in Na^+^ and K^+^ (membrane potential = -70 mV). We conducted of 16 replicas of unbiased MD simulation for an aggregated time of 7.3 µs for PF-NIS–ReO_4_^-^ and 6.8 µs for WT-NIS–ReO_4_^-^. Each replica was initialized with different Boltzmann-distributed velocities at 310.15 K in the isobaric-isothermal (NPT) statistical ensemble utilizing a V-rescale thermostat and a C-rescale barostat. PF-NIS was loaded with ReO₄⁻ and three bound Na^+^ ions initially placed at the sites observed by cryo-EM. We used the program GROMACS a simulation engine with a CHARMM36 force field modified to include non-bonded interactions with the oxyanion ReO_4_^-^ The system was gradually equilibrated using a six-step protocol suggested by CharMM-GUI ^55–57^.

## Data availability

Cryo-EM density maps and atomic coordinates have been deposited at https://drive.google.com/drive/folders/1nz0oJ7T8bpth-ZgGtew6J6MGmhOK9IjT?usp=sharing

## Author contributions

Alejandro Llorente-Esteban (A.LL-E.) and Nancy Carrasco (N.C.) conceived the project. A.LL-E. and Kendra Hoffsmith(K.H.) generated the plasmids with the amino acid substitutions at position 250. A.LL-E. generated the plasmids with the amino acid substitutions at positions 253 and 254. The plasmid containing PF-NIS was designed and generated by A.LL-E. The biochemical characterization of the position 250 mutants was carried out by A.LL-E., K.H., and Andrea Reyna-Neyra (A.R-N.); that of the position 253 and 254 mutants, by A.LL-E. and K.H.; and that of PF-NIS, by A.LL-E. and Haswitha Sabbineni (H.S.). The clonogenic survival experiments were performed by A.LL-E. and H.S. The electrophysiology experiments were designed and performed by Rían W. Manville (R.W.M.) and Geoffrey W. Abbott (G.W.A.). PF-NIS was purified, and the cryo-grids set up, by A.LL-E. and David Alejandro López-González (D.A.L-G.). The cryo-EM data set was analyzed by A.LL-E. The atomic model was fitted to the cryo-EM density map by A.LL-E. and Mario Antonio Bianchet (M.A.B.). The molecular dynamics simulations were carried out by M.A.B. and J. Alfonso Leyva. The manuscript was written by A.LL-E. and N.C., with input from all other authors.

## Acknowledgements

We are grateful to Bruno Lança for first drawing our attention to NIS residues L253 and V254, to Zenab Mchaourab for participating in the initial characterization of the G250 NIS mutants, to Paola Bisignano and Richard A. Stein for insightful discussions, and to the members of the Carrasco laboratory for critical reading of the manuscript. This study was supported by National Institutes of Health (NIH) grants #GM114250 (to N.C. and M.A.B.) and #GM130377 (to G.W.A.). We used the DORS storage system supported by the U.S. National Institute of Health (NIH) (S10RR031634 to Jarrod Smith). The molecular dynamics simulations were performed at the Advanced Research Computing at Hopkins (ARCH) core facility (rockfish.jhu.edu), which is supported by the National Science Foundation (NSF) under grant OAC1920103. Additional GPU resources were provided by DeltaAI and PSC Bridges-2 via allocation [BIO250233] through the Advanced Cyberinfrastructure Coordination Ecosystem: Services & Support (ACCESS) program, which is supported by NSF grants 2138259, 2138286, 2138307, 2137603, and 2138296.

## Additional information

**Correspondence and requests for materials** should be addressed to Nancy Carrasco.

---

**Extended Data Fig. 1:**
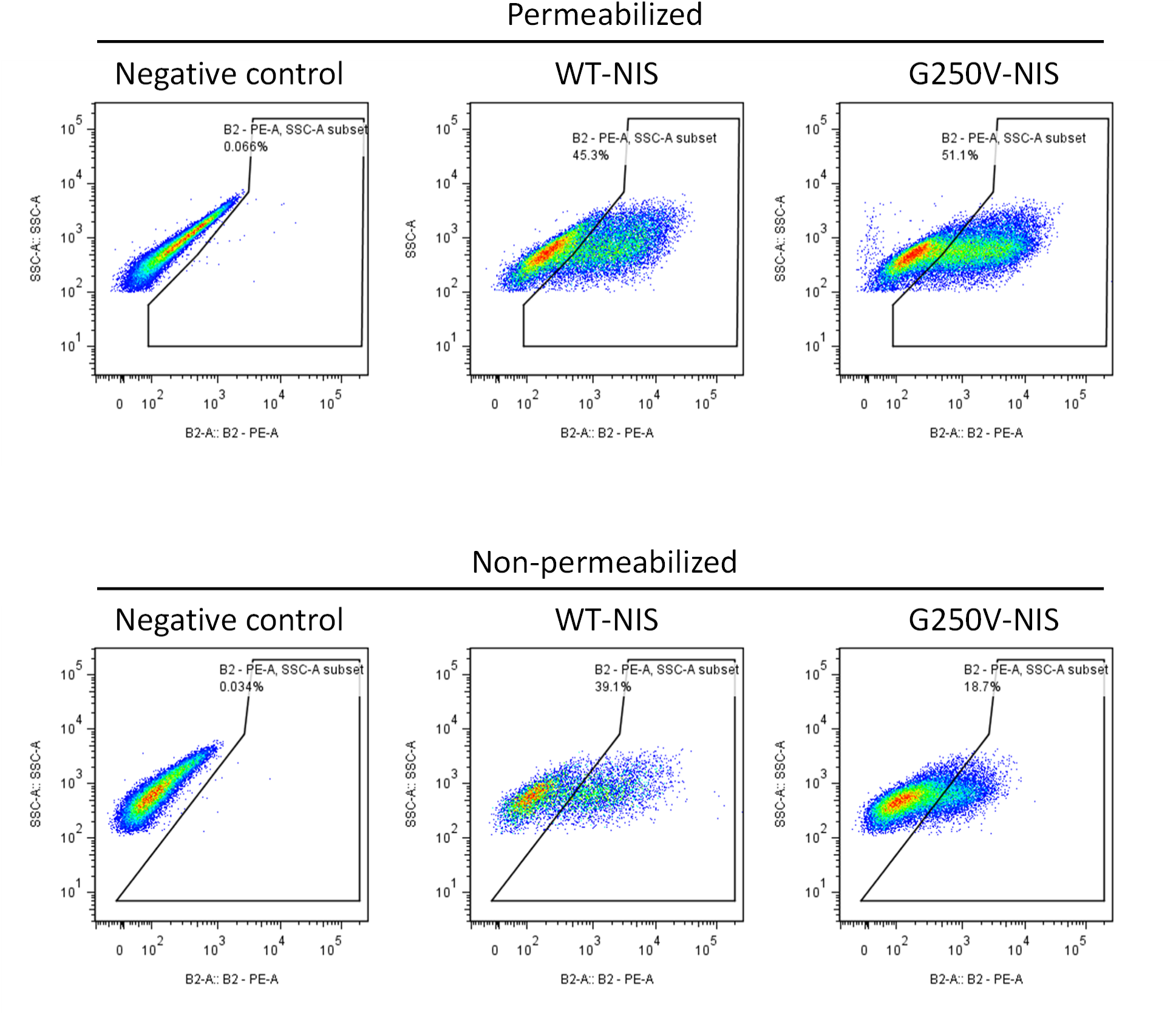
G250V-NIS is not targeted to the plasma membrane as robustly as WT-NIS. Cells were assayed by flow cytometry under permeabilizing and non-permeabilizing conditions to distinguish total and surface-expressed proteins, respectively. The gating strategy was established using non-NIS-expressing MDCK cells as a negative control for each condition. NIS-positive cells were labeled using a rat anti-HA IgG targeting an engineered HA-tag at the extracellularly facing N-terminus, followed by an anti-rat PE-conjugated secondary antibody. Forty-five percent of cells expressed WT-NIS under permeabilized conditions, whereas only 39% expressed it at the plasma membrane under non-permeabilized conditions, indicating that 86% of the WT-NIS molecules were localized at the plasma membrane. In contrast, G250V-NIS was detected in 39% of cells under permeabilized conditions, but only 19% of cells expressed it at the plasma membrane under non-permeabilized conditions, resulting in only 37% of the mutant NIS molecules reaching the plasma membrane. Data were analyzed and plotted using FlowJo version 7.6.2.

**Extended Data Fig. 2.**
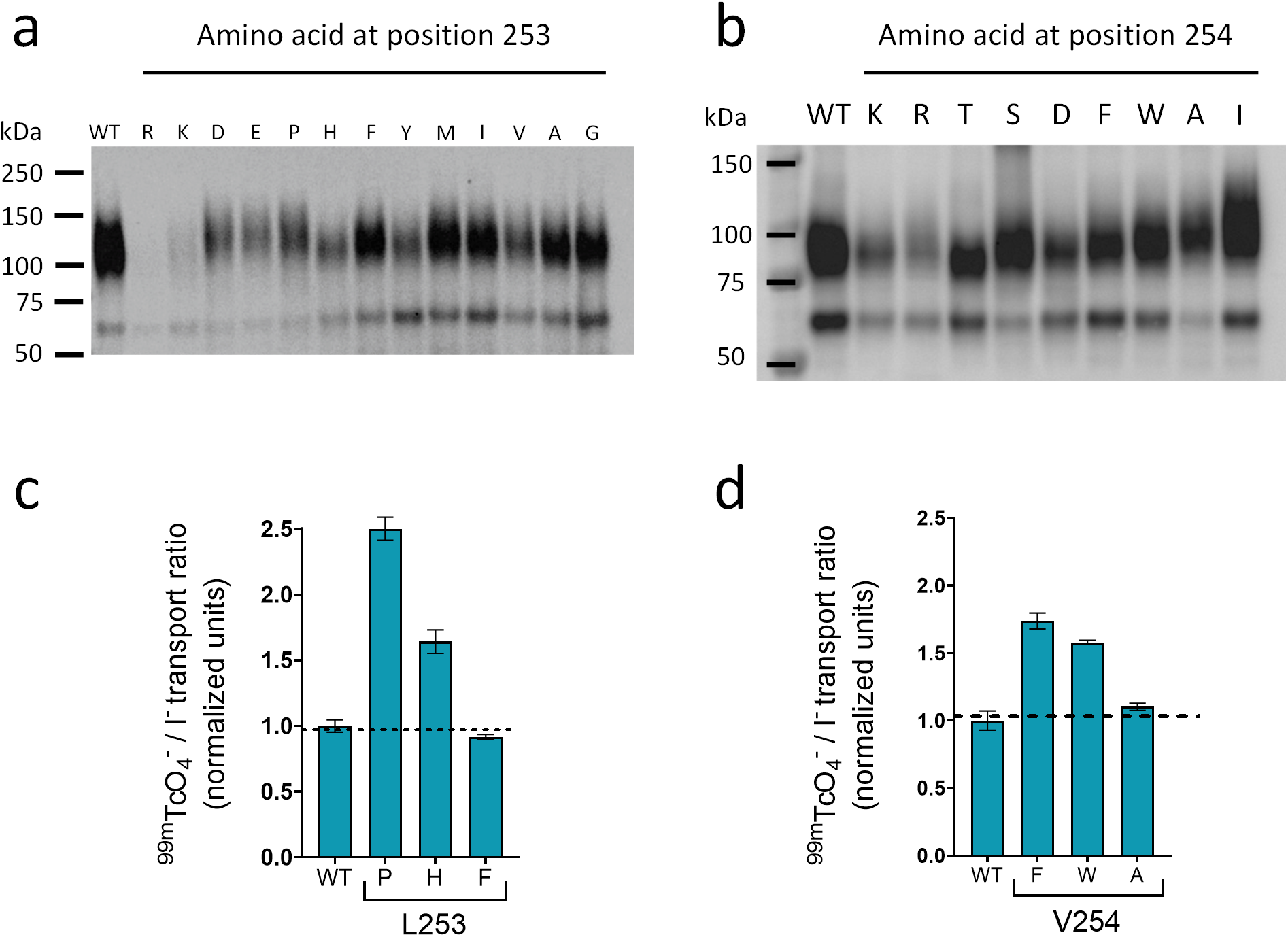
| Expression levels of mutant NIS proteins with substitutions at positions 253 and 254 that transport XO_4_^-^ better than I^-^. Western blot of MDCK cells expressing NIS mutants with amino acid substitutions at position **a,** 253 or **b,** 254. Membrane fractions (10 μg) were loaded for all samples. Ratios of ^99m^TcO_4_^-^ transport to I^-^ transport (normalized units) resulted from the analysis of the transport experiments shown in Fig. 2 b-e for selected amino acid substitutions at positions **c,** 253 and **d,** 254. Results are presented as mean ± SD.

**Extended Data Fig. 3.**
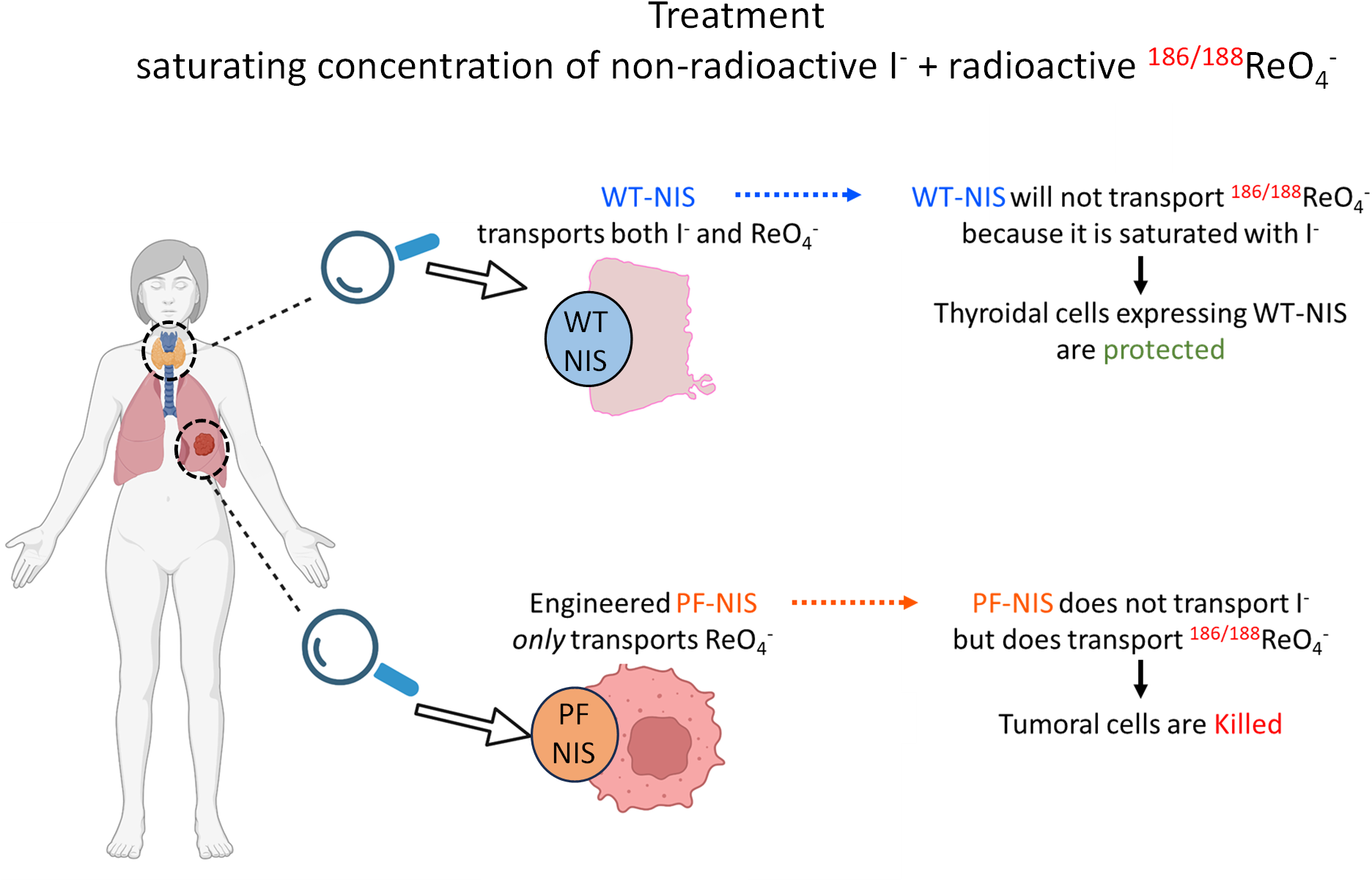
| Proposed therapeutic model using engineered NIS molecules transporting only ReO_4_^-^. An engineered NIS mutant that only transports XO_4_^-^s but not I^-^ is expressed in tumor cells (here, in the left lung) via gene therapy, using a tumor-specific promoter to restrict its expression. The patient then receives a saturating concentration of non-radioactive I^-^, along with radioactive ^186/188^ReO_4_^-^. Because the endogenous thyroidal WT-NIS is saturated by the non-radioactive I^-^, little or no ^186/188^ReO_4_^-^ enters the thyroid. By contrast, tumor cells expressing the engineered non-I^-^-transporting NIS (PF-NIS) continue to transport ^186/188^ReO_4_^-^, thereby targeting the therapeutic radioisotope to the tumor and killing the tumor cells while sparing the healthy thyroid. Figure created with BioRender.

**Extended Data Fig. 4.**
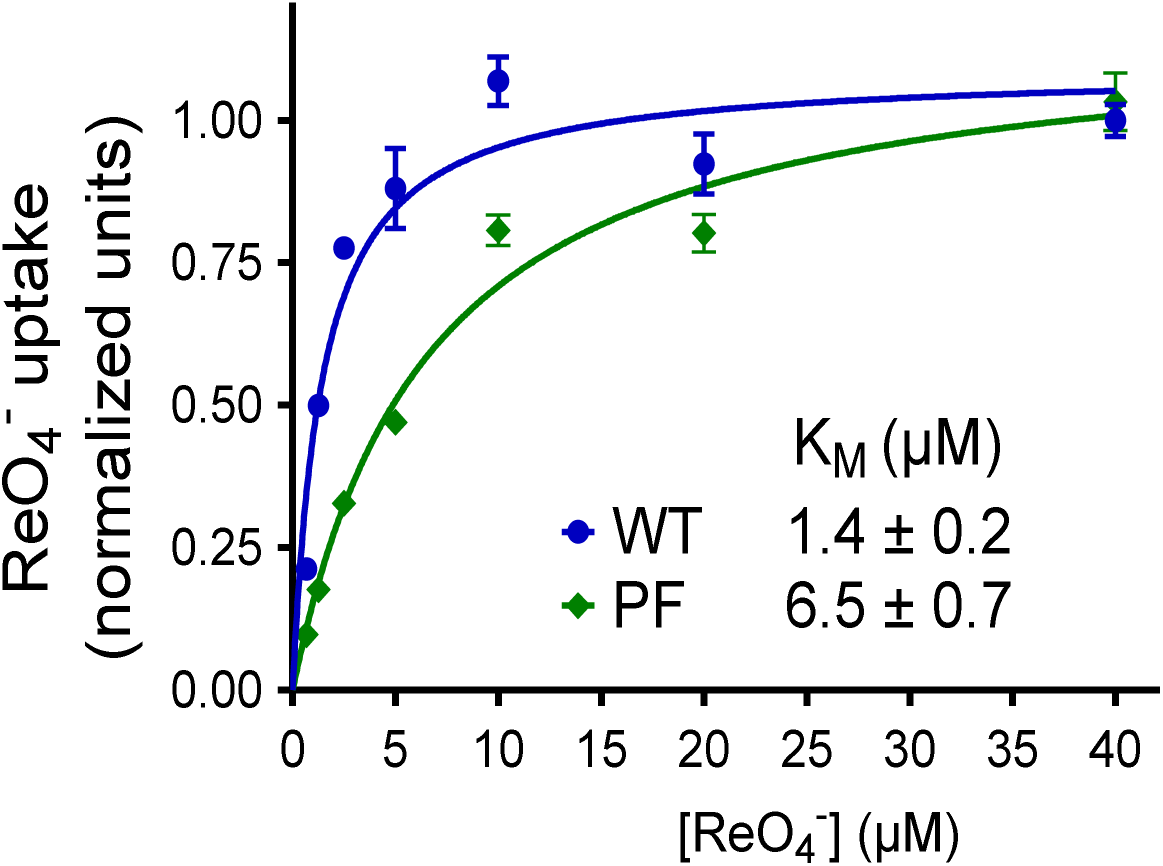
| PF-NIS has an only modestly lower apparent affinity for ReO_4_^-^ than WT-NIS. Initial rates of ReO_4_^-^ transport as a function of increasing concentrations of ReO_4_^-^ by WT-NIS and PF-NIS. Values are the mean ± s.e.m. of three biological replicates, each performed in triplicate (n ≥ 9).

**Extended Data Fig. 5.**
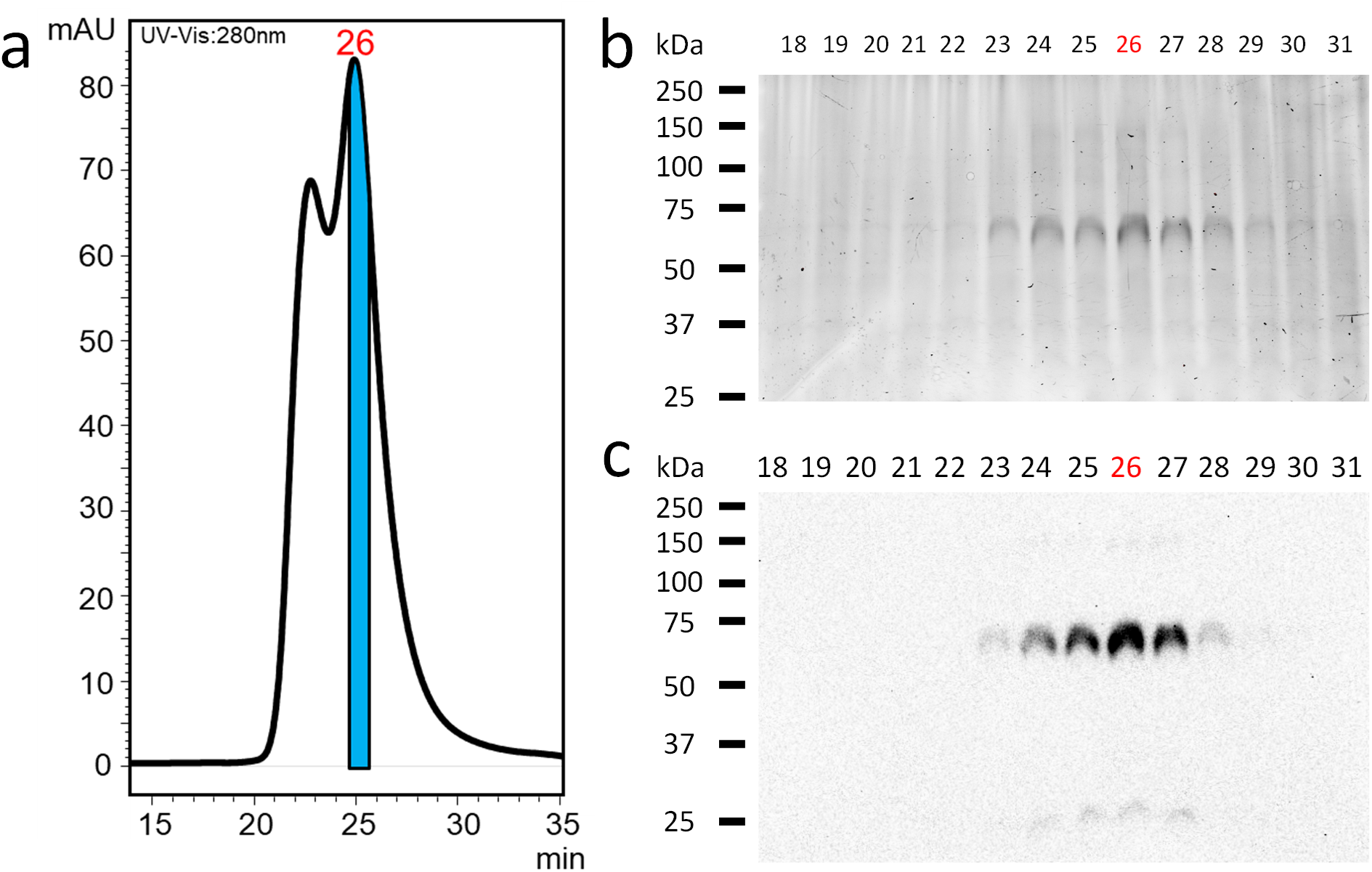
| Purification of PF-NIS. **a,** Size-exclusion chromatography (SEC) profile of PF-NIS, monitored by absorbance at 280 nm. The first peak represents protein aggregates unrelated to PF-NIS, while the second peak at fraction 26 corresponds to affinity-purified PF-NIS following two rounds of SEC on a Superdex 200 Increase column collecting at 1 fraction/minute. **b,** SDS–PAGE analysis of total protein on a Bio-Rad stain-free gel from each fraction corresponding to (a). Twenty µl out of 400 µl were loaded onto the gel from each fraction. **c,** Western blot of purified PF-NIS after transferring the gel shown in (b). The single band observed between ∼50 and ∼75 kDa in fractions 23–28 corresponds to non-glycosylated PF-NIS, with fraction 26 containing the highest amount of protein.

**Extended Data Fig. 6.**
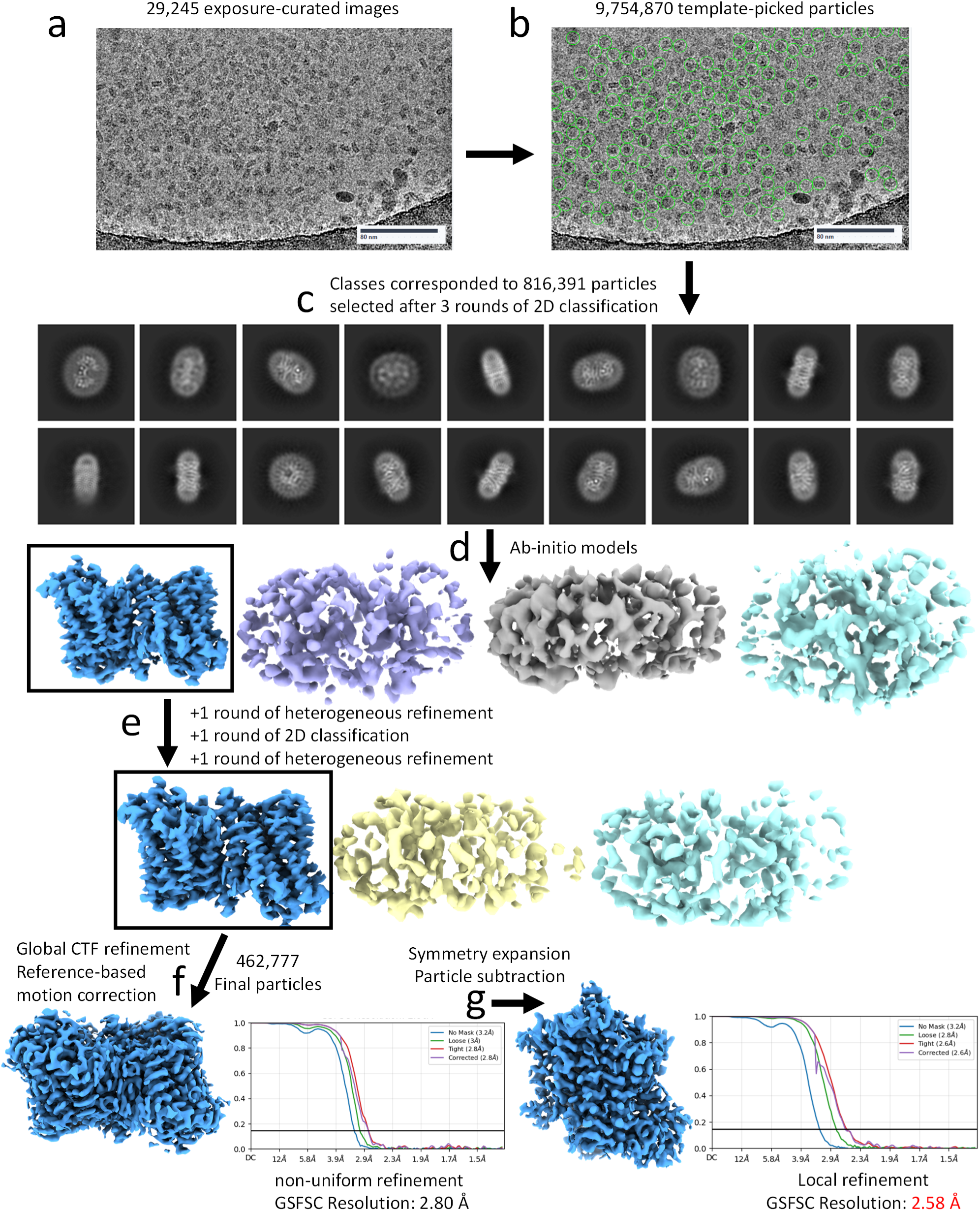
| Cryo-EM data processing and structural analysis of PF-NIS. **a,** Representative cryo-EM micrograph of PF-NIS from a dataset of 29,245 micrographs. **b,** Representative template-picked particles from the micrograph shown in (a). A total of 9,754,870 particles were selected from the entire dataset. **c,** Selected 2D class averages obtained after multiple rounds of 2D classification. **d,** Ab-initio models generated using CryoSPARC v4.5.3. **e,** Further classification of particles by heterogeneous refinements and additional 2D classification. **f,** A total of 462,777 particles were used for non-uniform refinement, yielding a 2.8 Å resolution map of the PF-NIS dimer. **g,** Symmetry expansion, particle subtraction, and local refinement of one monomer produced a final map at 2.58 Å resolution. Refined density maps and their Fourier shell correlation (FSC) curves are shown.

**Extended Data Fig. 7.**
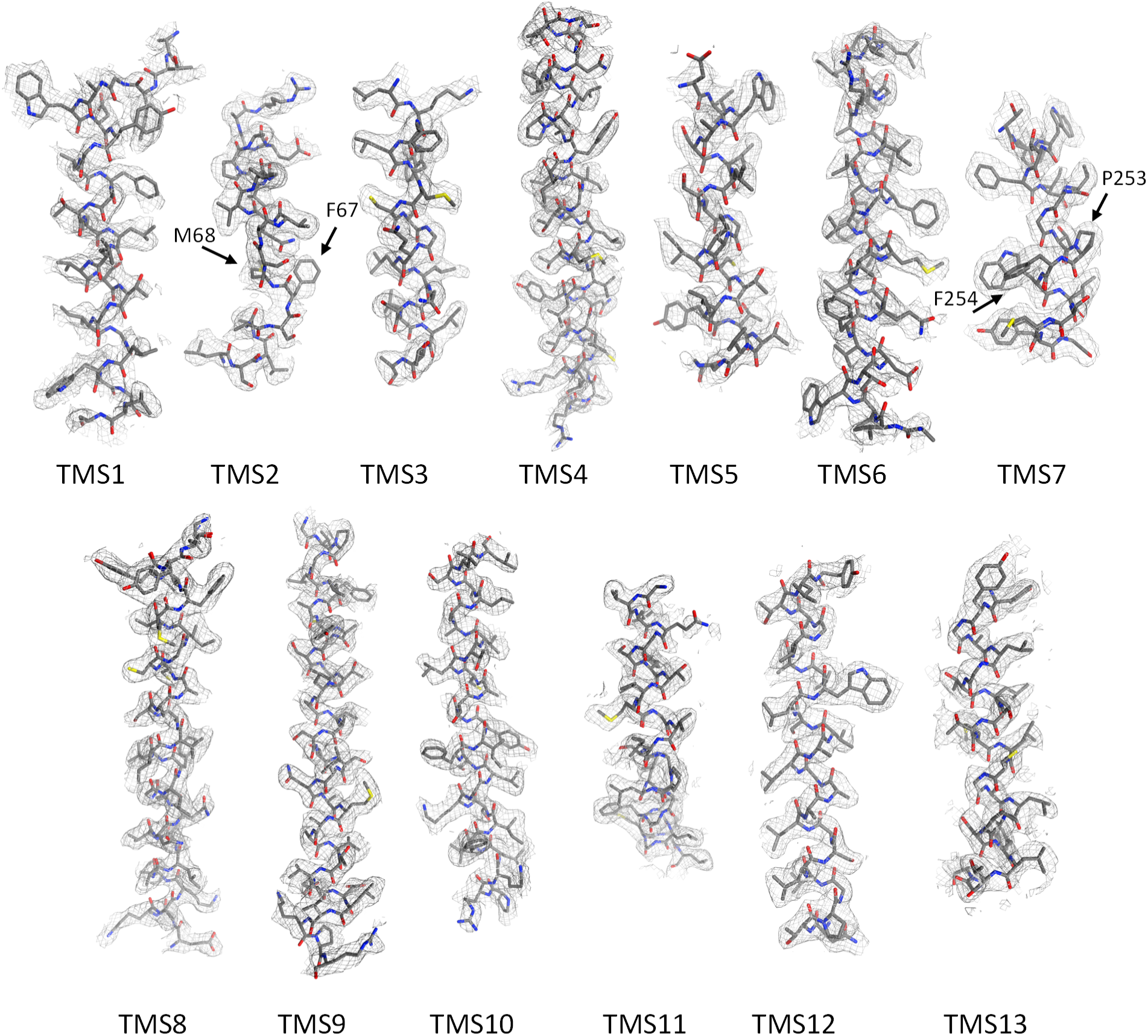
| Atomic model fitted into the cryo-EM density map. Cryo-EM density maps for all TMSs in PF-NIS, shown as black meshes. The fitted atomic model, depicted in grey, shows the protein backbone and side chains as sticks within the density. Sulfur, oxygen, and nitrogen atoms are color-coded as yellow, red, and dark blue, respectively. The residues F67, M68, P253, and F254 are indicated with black arrows.

**Extended Data Fig. 8.**
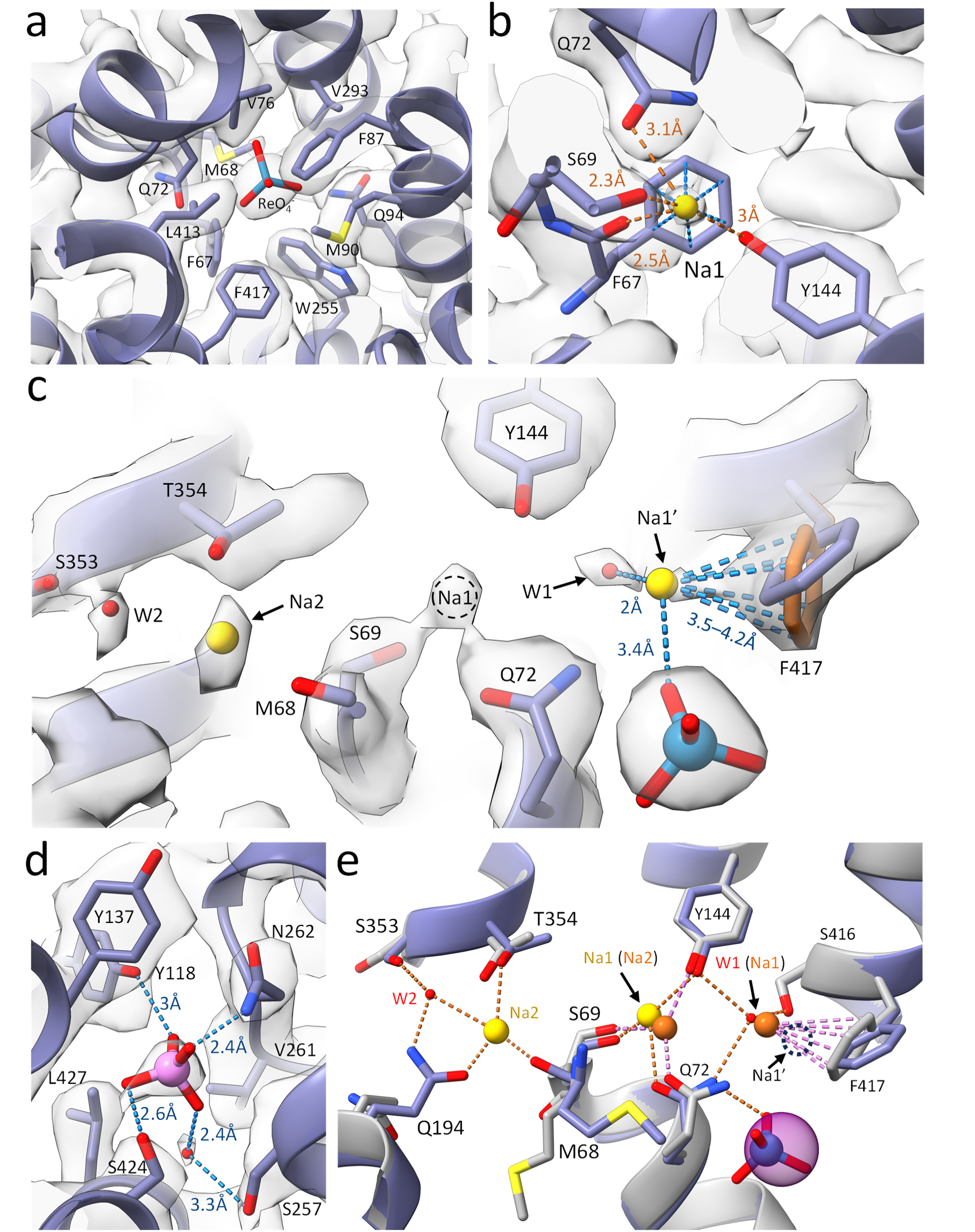
| Additional structural features of the PF-NIS structure. **a,** The ReO_4_^-^ binding site in PF-NIS. **b,** Na⁺ coordination at the Na1 site. F67 shows a plausible cation–π interaction with Na⁺ (all aromatic ring carbons are 4.8–5.4 Å from Na⁺, blue distances) and an additional coordination via its main-chain carbonyl at 2.5 Å (orange line), which are not observed in WT-NIS. **c,** Plausible alternative Na⁺ coordination at a Na1ʹ site at 3.4Å from ReO_4_^-^, involving a cation–π coordination with an alternative conformation of the F417 side chain suggested by its crescent shape at low thresholds (orange rotamer), and the water molecule at W1. The alternative rotamer of F417 positions all carbons of the aromatic ring at distances of 3.5–4.2 Å. This interpretation is only supported when the cryo-EM map is contoured at a very low threshold (5.7 rmsd), and was therefore considered insufficiently convincing for model deposition. The circular dotted line marks the primary position of Na⁺ as deposited in the atomic model at the Na1 site. **d,** Isolated tetrahedral density within a cavity formed by residues Y118, Y137, A140, S257, V261, N262, S424, and L427. For illustrative purposes, an oxyanion (pink) was fitted into the density, but it was not included in the final model due to uncertainty about its identity. **e,** Structural alignment between the determined structure of PF-NIS and the atomic model for WT-NIS-I^-^ (PDB: 7UV0). The densities assigned to Na⁺ ions in PF-NIS are shown in yellow, with their coordination depicted by orange dotted lines. The densities originally assigned Na⁺ ions in WT-NIS are shown in orange, with their coordination represented by pink dotted lines. The position of the hypothesized Na1ʹ site, suggested by the PF-NIS electron density map is indicated by a dotted circle. Red spheres indicate water molecules in the PF-NIS structure. I^-^ (semitransparent purple sphere) and ReO_4_^-^ (blue ball and red sticks) are shown at the anion binding site.

**Extended Data Fig. 9.**
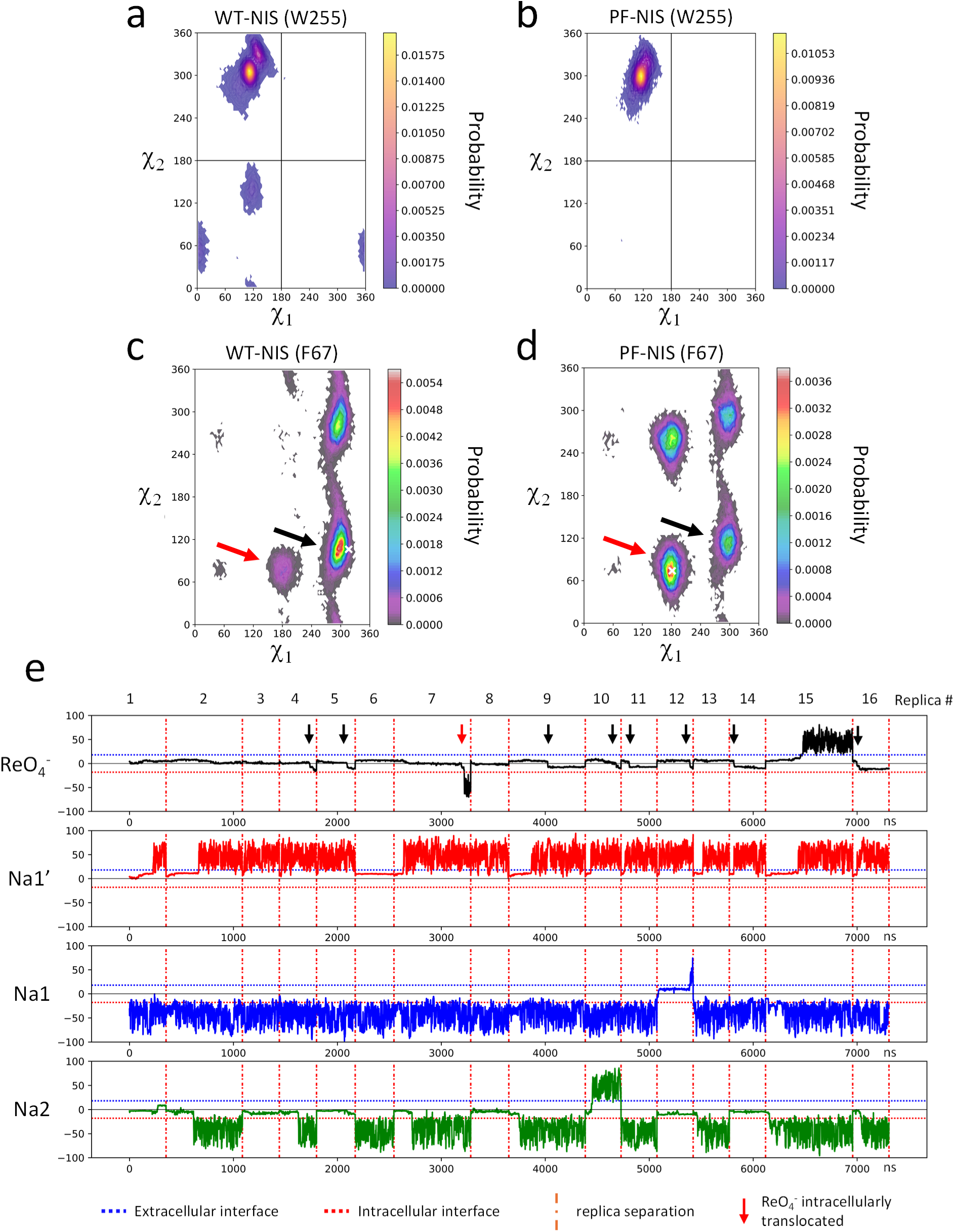
| Molecular dynamics simulations of PF-NIS–ReO_4_^-^. MD simulations show that the Janin (χ₁, χ₂) distributions for the side chain of W255 in **a,** WT-NIS explore a broader χ₁/χ₂ space than in **b,** PF-NIS. **c,** In WT-NIS, F67 more frequently adopts the *gauche*⁻ rotamer (χ_1_ ≈ 300°, χ₂ ≈ 100°; black arrow)—consistent with a lower-energy state—which facilitates the opening of the protein toward the cytosol more than does the less frequent *trans* rotamer (χ_1_ ≈ 180°, χ₂ ≈ 80°; red arrow) that occludes the pathway. **d,** PF-NIS favors the *trans* rotamer (red arrow), with the *gauche*⁻ rotamer (black arrow) being less frequently populated, consistent with a higher energetic penalty for the *gauche*⁻ rotamer. White X’s mark the positions of F67 in the experimentally determined structures. **e,** Aggregated 7.3 µs time series of the ions’ *z* coordinates for PF-NIS, representing transport along the translocation pathway. The *z* coordinate is centered at the membrane midplane, with zero marking the interface between the ion-binding pocket and the intracellular exit vestibule. Ions reaching negative *z* values appear committed to intracellular release, as no backward transitions to positive *z* values are observed. The cations at the Na1 and Na2 sites are transported and released intracellularly—except in replicas 12 and 10, respectively, where they escape. ReO_4_^-^ remains bound in most simulations, with only one escape event (replica 15) and one fully successful transport event (replica 7, red arrow). Additionally, in 9 out of 16 replicas (black arrows), ReO_4_^-^ progresses into the intracellular vestibule (*z* < 0) and is about to be transported. The cation at the Na1ʹ site escapes in all replicas.

**Extended Data Table 1.**

|  |  |
| --- | --- |
|  | PF-NIS<br>map and model available at:<br>https://drive.google.com/drive/folders/1nz0oJ7T8bpth-ZgGteW6J6MGmhOK9IjT?usp=sharing |
| **Data collection and processing** |  |
| Magnification | 130,000 x |
| Voltage (kV) | 300 |
| Electron exposure (e-/Å<sup>2</sup>) | 55 |
| Defocus range (μm) | -2.3 to -0.8 |
| Pixel size (Å) | 0.647 |
| Symmetry imposed | C1 |
| Initial particle images (no.) | 9,754,870 |
| Final particle images (no.) | 462,777 |
| Map resolution (Å) | 2.58 |
| FSC threshold | 0.143 |
| Map resolution range (Å) | 2.47 to 3.594 |
| **Refinement** |  |
| Initial model used (PDB code) | 7UUZ |
| Model resolution (Å) | 2.6 |
| FSC threshold | 0.143 |
| Model resolution range (Å) | 2.4-2.9 |
| Map sharpening *B* factor (Å<sup>2</sup>) | 128.8 |
| Model composition |  |
| Non-hydrogen atoms | 3812 |
| Protein residues | 506 |
| Ligands | 2Na<sup>+</sup> & 1 ReO<sub>4</sub><sup>-</sup> |
| *B* factors (Å<sup>2</sup>) |  |
| Protein | 37.6 |
| Ligand | 71.8 |
| R.m.s. deviations |  |
| Bond lengths (Å) | 0.004 |
| Bond angles (°) | 0.625 |
| Validation |  |
| MolProbity score | 2.62 |
| Clashscore | 11.31 |
| Poor rotamers (%) | 7.79 |
| Ramachandran plot |  |
| Favored (%) | 94.6 |
| Allowed (%) | 5.2 |
| Disallowed (%) | 0.2 |

